# Catalytic-dependent and independent functions of the histone acetyltransferase CBP promote pioneer factor-mediated zygotic genome activation

**DOI:** 10.1101/2024.10.04.616638

**Authors:** Audrey J. Marsh, Sergei Pirogov, Abby J. Ruffridge, Suresh Sajwan, Tyler J. Gibson, George Hunt, Yadwinder Kaur, Melissa M. Harrison, Mattias Mannervik

## Abstract

Immediately after fertilization the genome is transcriptionally quiescent. Maternally encoded pioneer transcription factors reprogram the chromatin state and facilitate the transcription of the zygotic genome. In *Drosophila*, transcription is initiated by the pioneer factor Zelda. While Zelda-occupied sites are enriched with histone acetylation, a post-translational mark associated with active *cis*-regulatory regions, the functional relationship between Zelda and histone acetylation in zygotic genome activation remained unclear. We show that Zelda-mediated recruitment of the histone acetyltransferase CBP is essential for zygotic transcription. CBP catalytic activity is necessary for release of RNA Polymerase II (Pol II) into transcription elongation and for embryonic development. However, CBP also activates zygotic transcription independent of acetylation through Pol II recruitment. Neither acetylation nor CBP are required for the pioneering function of Zelda. Our data suggest that pioneer factor-mediated recruitment of CBP is a conserved mechanism required to activate zygotic transcription but that this role is separable from the function of pioneer factors in restructuring chromatin accessibility.

## Introduction

Immediately following fertilization, the newly formed genome is transcriptionally silent. This allows for the zygotic genome to be rapidly reprogrammed to enable the generation of a new, unique organism. This fast-paced and conserved period of metazoan development is called the maternal-to-zygotic transition (MZT), in which maternal mRNAs and proteins loaded into the egg during oogenesis trigger expression of the zygotic genome after fertilization^1^.

Across animals, reprogramming during the MZT is driven by pioneer factors. In contrast to many transcription factors for which nucleosomes are a barrier to binding, pioneer factors bind nucleosomes and reorganize chromatin accessibility. These changes to chromatin accessibility result in the recruitment of downstream transcription factors and the activation of new gene expression programs. As a result, pioneer factors act at the top of transcriptional networks to drive developmental transitions^2,3^. Understanding pioneer factor function is therefore critical for elucidating how the genome is interpreted through dynamic developmental transitions.

The first major activator of zygotic transcription, Zelda (Zld), was initially identified in *Drosophila melanogaster*^4^, and *Drosophila* have continued to be a powerful model for studying the regulatory principles that govern how pioneer factors remodel the genome and drive development. In *Drosophila*, the MZT occurs over the first three hours after egg laying (AEL). During this period, the nuclei undergo 14 nuclear division cycles (NC) within a syncytium^5^. At NC8, maternally encoded *zld* is translated leading to the initiation of zygotic transcription^6–8^. Transcription from the zygotic genome is gradually activated with widespread transcription occurring at NC14, coincident with the slowing of the division cycle. Zld is required for chromatin accessibility at hundreds of *cis-*regulatory elements^9,10^. This accessibility potentiates the binding of additional transcription factors and promotes proximal gene expression^4,9–14^. Zld binds to nucleosomes *in vitro*, and ectopic expression of Zld in culture induces chromatin accessibility^15–17^. Together these studies demonstrate that Zld is a pioneer factor necessary for reprogramming the zygotic genome. Nonetheless, how Zld facilitates chromatin accessibility after nucleosome binding remains unclear. Here, we explore mechanisms of Zld pioneer activity to understand how pioneer factors facilitate dramatic changes in transcription during developmental transitions.

Prior to Zld-mediated genome activation, the histone tails are largely devoid of post-translational modifications as assayed by chromatin immunoprecipitation followed by sequencing (ChIP-seq)^18^. Acetylation of H3K18, H3K27, and H4K8 is detected initially at NC8 and is enriched at Zld-bound regions^9,18,19^. Similarly, in zebrafish H3K27ac accumulates prior to activation of the zygotic genome, and pioneer factors are required for the deposition of this mark^20–22^. While acetylation is thought to promote chromatin accessibility by neutralizing the positive charges of lysine groups on histone tails, the connection between histone acetylation, gene expression and chromatin accessibility remains largely correlative^23,24^. Histone acetylation also recruits bromodomain containing nucleosome remodelers to clear nucleosomes from *cis-*regulatory regions^25^. Thus, Zld may promote chromatin accessibility through the recruitment of a histone acetyltransferase (HAT).

Acetylation of H3K18, H3K27 and H4K8 are dependent upon the deeply conserved CREB-binding protein (CBP)/p300 HAT family^26–32^. CBP/p300 is a widespread transcriptional coactivator. Indeed, CBP/p300 binding and the acetylation of H3K27 it catalyzes are markers used to identify active *cis-*regulatory regions^33,34^. The catalytic core of CBP/p300 is functionally required for HAT activity and includes a bromodomain, RING, PhD finger, HAT, ZZ, and TAZ domains. Catalytic activity must be activated through acetylation of an autoinhibitory loop within the HAT domain^35–37^. Therefore, CBP/p300 occupancy alone is not predictive of its catalytic activity or whether CBP-bound sites are enriched with histone acetylation^38^. In fact, non-catalytic activities of CBP/p300 contribute to the regulation of gene expression^30,39^. Because *Drosophila* have a single CBP/p300 homologue, encoded by the gene *nejire,* they provide a simplified system to investigate the possible relationship between CBP/p300 and pioneer factors in the early embryo^40^.

Here, we investigated the connection between Zld pioneer activity, acetylation, and gene expression. We discovered that Zld is required for CBP recruitment to the genome at a subset of sites. CBP mediates zygotic gene expression and is necessary for embryonic development. CBP activates gene expression through two distinct mechanisms that depend on catalytic-dependent and independent activities. Independent of catalytic activity, CBP is needed for the recruitment or stability of RNA polymerase II (Pol II) at gene promoters. Pol II initiates but pauses 40-60 bp downstream of the transcription start site at many zygotic genes^41^. Release from this promoter-proximal pausing depends on the kinase activity of P-TEFb^42,43^. We found that the catalytic activity of CBP is required for Pol II pause release and robust transcription. In contrast to the essential role of CBP in zygotic gene expression, CBP is dispensable for chromatin accessibility. Thus, Zld pioneer activity is independent of its ability to recruit CBP and is separable from activating gene expression. These data suggest that together Zld and CBP coordinate the activation of the zygotic genome and elucidate a mechanism by which pioneer factors recruit cofactors to reprogram transcription and drive developmental transitions.

## Results

### Zld is required for recruitment of CBP to a subset of loci

The correlation between Zld-bound sites and CBP-dependent histone acetylation suggested that Zld might recruit CBP to these sites to catalyze acetylation. To confirm CBP is expressed during the MZT, we endogenously tagged the N-terminus of CBP with GFP using Cas9-mediated genome engineering (Figure S1A). CBP^GFP^ was evident in nuclei throughout the MZT beginning as early as NC10 and into NC14 (Figure 1A). ChIP-seq for CBP^GFP^ on hand-sorted stage 5 (NC14) embryos identified 5,479 regions bound by CBP. To avoid technical differences in peak numbers, we used the same ChIP-seq protocol to identify Zld-bound loci at stage 5, when the major wave of zygotic transcription initiates. Based on the overlap of Zld-bound regions with CBP-dependent histone acetylation marks, we predicted that CBP and Zld would have overlapping genome occupancy. Indeed, we identified 2,932 Zld-bound sites, of which 1202 were shared with CBP (Figure 1B). For downstream analysis, we split the total combined Zld and CBP^GFP^ peaks into three classes: shared, Zld-unique, and CBP-unique. (Figure 1C-E). Annotation of individual peaks by the type of *cis-*regulatory element showed that CBP occupancy is largely at promoters, as both shared (52.7%) and CBP unique (67.0%) classes have a high proportion of promoter-bound sites as compared to other genomic elements. (Figure 1F).

**Figure 1.**
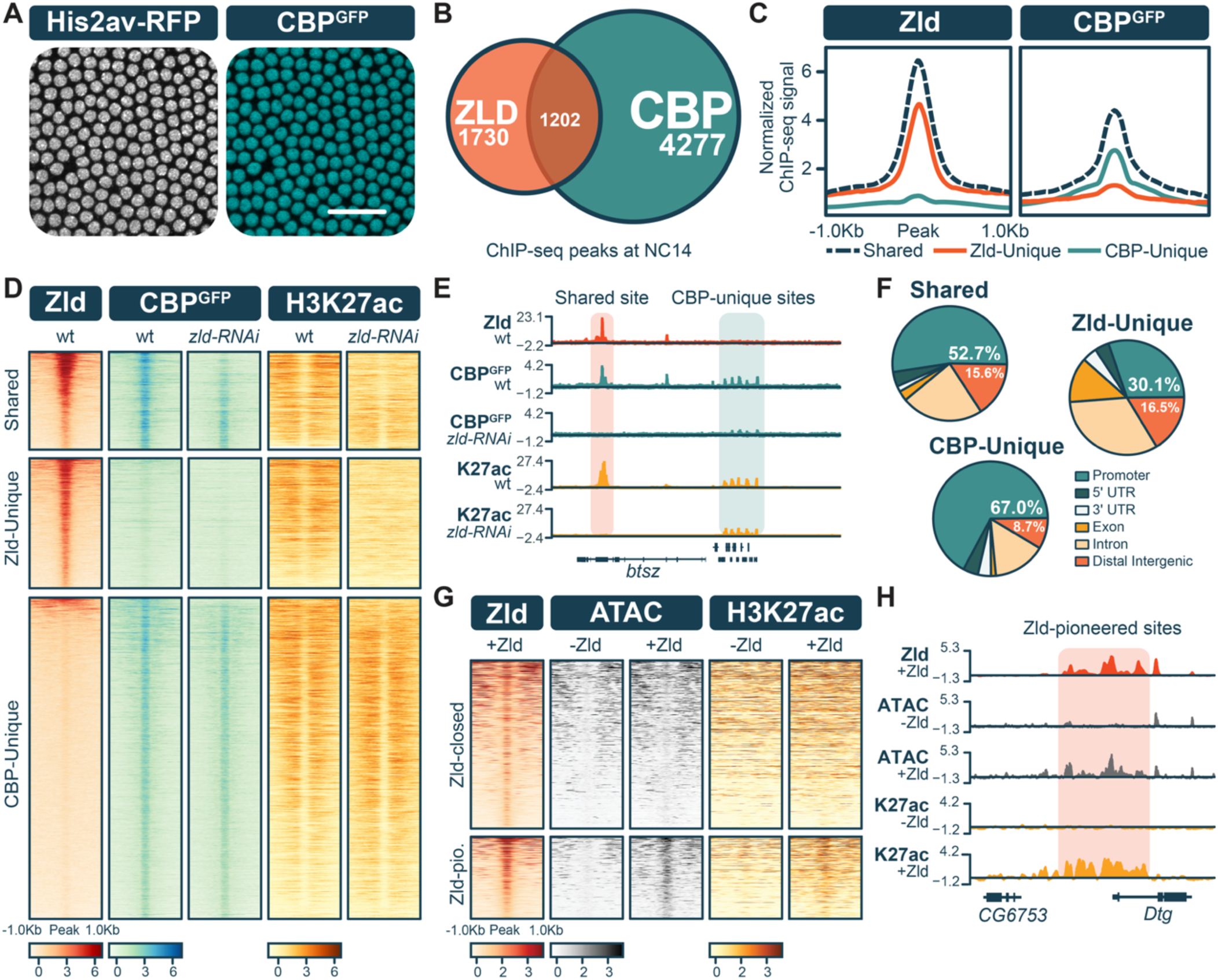
The pioneer factor Zld is required for CBP recruitment and H3K27ac deposition at co-occupied sites and is sufficient to induce H3K27ac at Zld-pioneered regions. **(A)** His2AvRFP (white) or CBP^GFP^ (teal) at NC14 (stage 5) in a *CBP^GFP^;His2AvRFP* embryo. Scale bar, 25µM. **(B)** Venn diagram showing the proportion of Zld and CBP^GFP^ ChIP-seq peaks that overlap in stage 5 embryos. **(C)** Metaplots of the Zld or CBP^GFP^ ChIP-seq signals centered on the peak. Peaks from each ChIP-seq experiment were divided into shared (dashed), Zld-unique (red), or CBP-unique (teal) classes. Data is normalized to background. **(D)** Heatmaps showing Zld, CBP^GFP^, or H3K27ac occupancy as measured by ChIP-seq in stage 5 wild-type (wt) or *zld-RNAi* embryos. All peaks are ranked by Zld-binding intensity. Zld and H3K27ac experiments are z-score normalized. CBP^GFP^ experiments are spike-in normalized. **(E)** Representative genome browser track of the *bitesize (btsz)* locus depicting peaks from the shared class (red bar) and CBP-unique class (teal bar). **(F)** Pie charts depicting the proportion of ChIP-seq peaks bound to distinct *cis-*regulatory elements. Promoters were defined as -500bp and +100bp from the transcription start site (TSS) of the closest coding region to each peak. **(G)** Heatmaps of Zld occupancy (ChIP-seq), chromatin accessibility (ATAC-seq), and H3K27ac (CUT&RUN) at regions of closed chromatin bound by Zld upon expression in S2 cells. Peaks are separated based on whether they gain accessibility following Zld induction: Zld-closed (remain inaccessible) and Zld-pioneered (gain accessibility). Signals are z-score normalized. **(H)** Representative genome browser track from 1G, highlighting a region from the Zld-pioneered class (red bar) at the *Dpp target gene* (*Dtg*) locus.

CBP is incapable of binding directly to DNA. The overlap of Zld- and CBP-bound regions suggested that Zld might recruit CBP to co-occupied sites. To test this, we used RNAi to deplete maternally encoded Zld in CBP^GFP^ embryos and identified CBP-binding sites using ChIP-seq (Figure S1B)^10^. CBP^GFP^ binding was strongly reduced at shared regions but was maintained at loci corresponding to the CBP-unique peaks (Figure 1D,E). Thus, Zld is required to recruit CBP to the genome at shared regions. As expected, CBP-catalyzed H3K27ac was enriched at CBP-bound regions in wild-type embryos. Levels of H3K27ac were strongly reduced at Zld-bound sites in *zld-RNAi,* reflecting the loss of CBP occupancy (Figure 1D,E). Unexpectedly, H3K27ac was also reduced at the Zld-unique peaks, suggesting CBP may be dynamically recruited to these regions in a manner that is not captured by ChIP-seq. This is supported by evidence that H3K27ac is exclusively dependent upon CBP in the early embryo, making it unlikely that another HAT is responsible for H3K27ac at these sites^30^. Thus, Zld is recruiting CBP to thousands of genomic loci and promoting H3K27ac.

From these data, it is unclear whether Zld is directly recruiting CBP through a protein-protein interaction or indirectly through promoting chromatin occupancy of another transcription factor. As expected, variations of the CAGGTAG motif to which Zld binds were the most enriched motifs at shared and Zld-unique classes. The top motifs identified in CBP-unique peaks are bound by the insulator proteins BEAF-32 and Dref, highlighting a potential distinct function of these regions compared to Zld-CBP shared sites (Figure S1C). Bicoid (Bcd), Dorsal (Dl), and Twist (Twi) are transcription factors that depend on Zld-mediated pioneer activity to access their binding motifs^11–13^. Therefore, these factors could function to indirectly recruit CBP to Zld-pioneered regions. The Dl motif was found to be enriched in all three peak classes, suggesting that at both Zld-bound and CBP-unique sites Dl might facilitate CBP binding. The Bcd motif was specific to Zld-bound classes, while the Twi motif was enriched in CBP-unique sites (Figure S1C). Analysis of published ChIP-seq data showed that binding of both Bcd and Dl at shared sites, but not CBP-unique sites, depended on Zld (Figure S1D,E)^10,13^. This further supports a possible role for both transcription factors in facilitating CBP binding at these loci. This analysis highlights factors that might function with Zld to direct CBP binding in the early embryo.

We showed that Zld is necessary for CBP binding to embryonic chromatin. To determine if Zld was sufficient to promote CBP recruitment, we exogenously expressed Zld in Schneider 2 cells (S2), where endogenous Zld expression is below the level of detection^16^. We previously used this system to show that exogenously expressed Zld bound closed chromatin and promoted accessibility at a subset of loci (Zld-pioneered sites)^16^. To investigate how Zld binding affects histone acetylation, we performed CUT&RUN for H3K27ac with and without Zld expression. We identified enrichment of H3K27ac specifically at the Zld-pioneered class upon Zld induction (Figure 1G,H and S2). Together with our data from embryos, this demonstrates that Zld is not only necessary for histone acetylation but is also sufficient for promoting this histone modification. While motif analysis in embryos suggested the possible involvement of other factors, such as Dl, Bcd and Twi, these factors are not expressed in S2 cells. Thus, Zld is either directly recruiting CBP to promote acetylation of closed chromatin in S2 cells or Zld-mediated accessibility facilitates the binding of other factors that promote CBP recruitment.

### Maternally encoded CBP is essential for zygotic genome activation

The correlation between Zld binding and CBP occupancy suggested that CBP might be essential for transcriptional activation during ZGA. We explored this hypothesis by depleting CBP^GFP^ protein using a maternally driven *deGradFP* transgene^44,45^. This construct expresses a GFP nanobody fused to an F-box-containing protein that can intercalate with an endogenous ubiquitin ligase complex to selectively degrade GFP-tagged proteins. We confirmed that this system results in depletion of CBP^GFP^ at NC14 (Figure 2A) and a robust decrease in H3K27ac and H3K18ac levels (Figure S3A,B). Hatching rates performed roughly 24h after egg laying (AEL) demonstrated that CBP^deGrad^ embryos were inviable, similar to zygotic mutants (Figure 2B)^40^. Comparable results were obtained with RNAi knockdown of *CBP*^30,46^. We conclude that maternally encoded CBP is essential for early embryo development.

**Figure 2.**
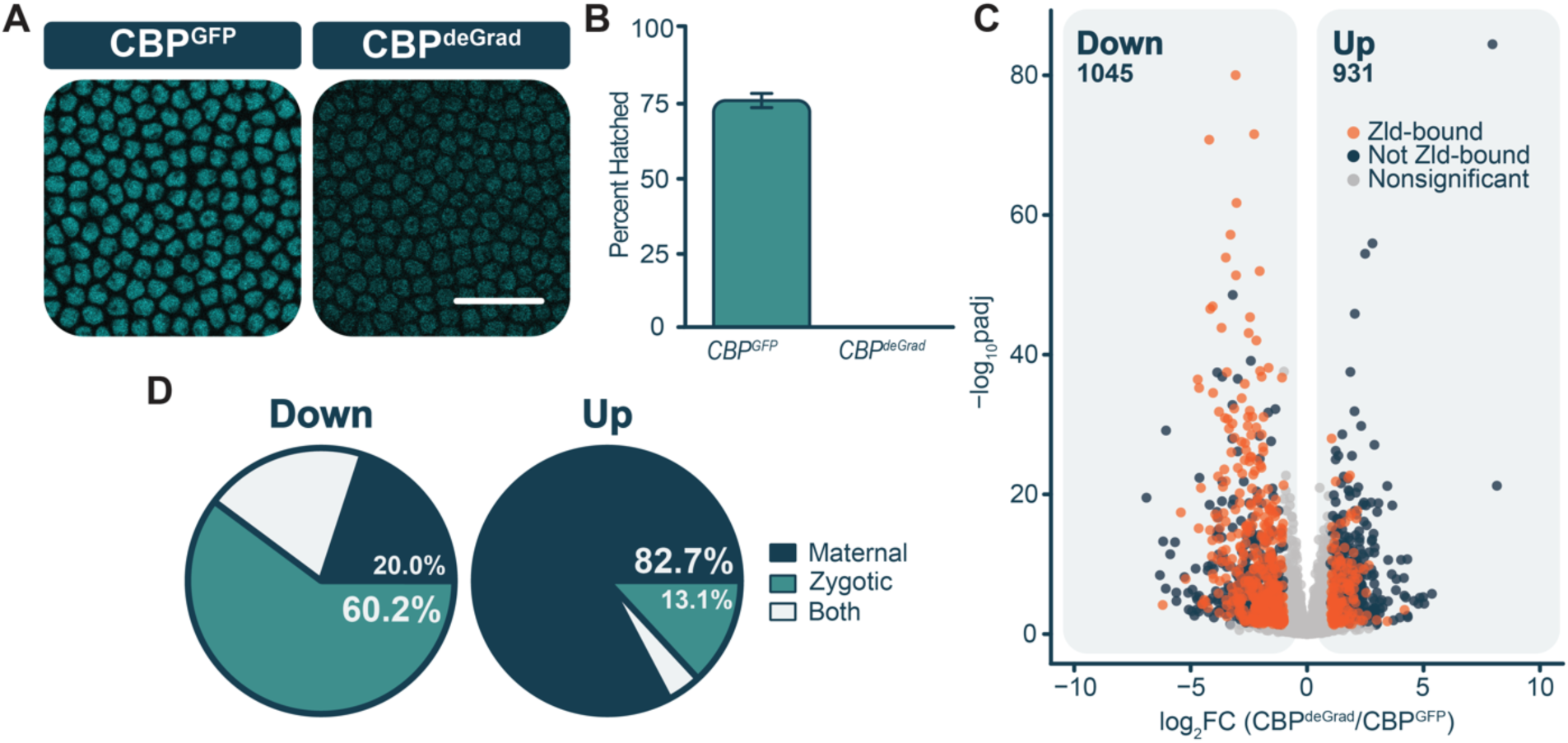
Maternal CBP is required for zygotic genome activation. **(A)** GFP signal in CBP^GFP^ control and CBP^deGrad^ embryos staged at NC14. Scale bar, 25uM. **(B)** Percent CBP^GFP^ (n= 388) or CBP^deGrad^ (n= 451) embryos hatched 24hrs AEL. Error bars are the standard deviation between three replicates. **(C)** Volcano plot of single embryo RNA-seq from CBP^deGrad^ embryos as compared to CBP^GFP^ embryos. Genes with Zld-proximal peaks are indicated in red. Navy represents genes that change in expression lacking a proximal Zld-binding sites, and grey represents genes with statistically insignificant changes in expression (adjusted p-value < 0.05, |log2 fold change| > 1). **(D)** Pie charts of the expression patterns of genes with decreased (left) or increased (right) expression in CBP^deGrad^ embryos as compared to CBP^GFP^ embryos during the MZT: maternal, zygotic, or both.

Using this tool to robustly deplete CBP from the embryo, we performed single-embryo mRNA-seq on control (*CBP^GFP^; His2AvRFP*) and *CBP^deGrad^;His2AvRFP* embryos harvested precisely 15 min into NC14. Differential analysis revealed 1045 genes with decreased expression upon CBP degradation and 931 with increased expression (Figure 2C). These genes are similarly dysregulated when CBP is depleted by RNAi (Figure S3C)^30^. Increased and decreased genes were classified as maternal or zygotically expressed based on prior analysis^47^. 337 increased and 485 decreased genes could not be classified, as they were not present within the original referenced dataset. Of the genes that were classifiable, 82.7% (491/594) of increased genes were maternally expressed, suggesting most of the increased genes are maternally provided mRNAs that fail to be efficiently degraded when CBP is absent. These are likely indirect targets of embryonically expressed CBP. By contrast, genes with decreased mRNA levels are enriched for those that are zygotically expressed (60.2%, 448/560 genes), and are therefore likely to be enriched for those directly activated by CBP (Figure 2D). 43% of these zygotic genes have CBP bound to a proximal region as assayed by ChIP-seq. This is likely an underestimate of CBP occupancy as ChIP-seq only reflects binding sites that enable cross-linking of CBP to chromatin. Thus, CBP binding functions broadly to activate gene expression during the MZT.

Given that CBP depends on Zld for recruitment to the genome at thousands of loci, genes that depend upon CBP for expression may be Zld targets. We therefore used immunoblots to test whether the effects of CBP depletion on gene expression were the indirect result of changes to Zld levels. Zld levels did not change in maternally depleted CBP embryos (Figure S3D). Because Zld binds and activates hundreds of zygotically expressed genes, we would expect that Zld-binding sites would be enriched near those genes that depend on CBP for expression. Indeed, 37.9% (396/1045) of decreased genes had a proximal Zld-binding site, which was more than 2.5x higher than the enrichment near increased genes (14.2% (132/931)) (Figure 2C). To further test the connection between CBP- and Zld-mediated gene expression, we analyzed single-embryo RNA-seq from embryos in which maternal Zld was optogenetically inactivated during zygotic genome activation^15^. The log_2_ fold change in expression of Zld-dependent genes correlated with the log_2_ fold change of the same genes in CBP^deGrad^ RNA-seq. Genes that were decreased in Zld-inactivated embryos were also down in CBP^deGrad^, indicating that Zld-dependent genes also require CBP for activation (Figure S3E). These results cumulatively support a model in which Zld recruitment of CBP is necessary to mediate gene expression during the MZT.

### CBP-mediated acetylation is required in the zygote for embryonic development

Both the previously published RNAi knockdown and our deGrad depletion rely on expression in the maternal germline^30,45^, making it impossible to disentangle effects caused by any knockdown in the germline as compared to the early embryo. To overcome this challenge, we used an optogenetic strategy that has previously been used to precisely inactivate transcriptional activators during the period of zygotic expression (NC10-14) and, in so doing, circumvents phenotypes nonspecific to ZGA that can arise when maternally expressed factors, like CBP, are attenuated during oogenesis^15,48^. For this purpose, the blue-light responsive CRY2 polypeptide was engineered onto the N-terminus of endogenous CBP (Figure 3A). Inactivation of CBP^CRY2^ by blue light was tested by assaying for CBP-mediated histone acetylation levels (H3K27ac and H3K18ac) by immunoblot on extract from either wild-type or CBP^CRY2^ embryos treated with blue light from 1-3h AEL. CBP-mediated acetylation of H3K27 and H3K18 were robustly decreased in CBP^CRY2^ embryos exposed to blue light as compared to both wild-type controls or CBP^CRY2^ embryos kept in the dark (Figure S4A). CUT&Tag for histone acetylation on stage 5 embryos treated with blue light from 1-3 hr AEL showed a genome-wide decrease in H3K27ac and H3K18ac upon blue-light inactivation (Figure 3B). Blue-light treatment did not generally affect histone acetylation as the CBP-independent mark H3K9ac was unaffected after blue-light treatment (Figure 3B). These assays demonstrate that blue-light treatment during the MZT resulted in inactivation of the acetyltransferase activity of CBP^CRY2^.

**Figure 3.**
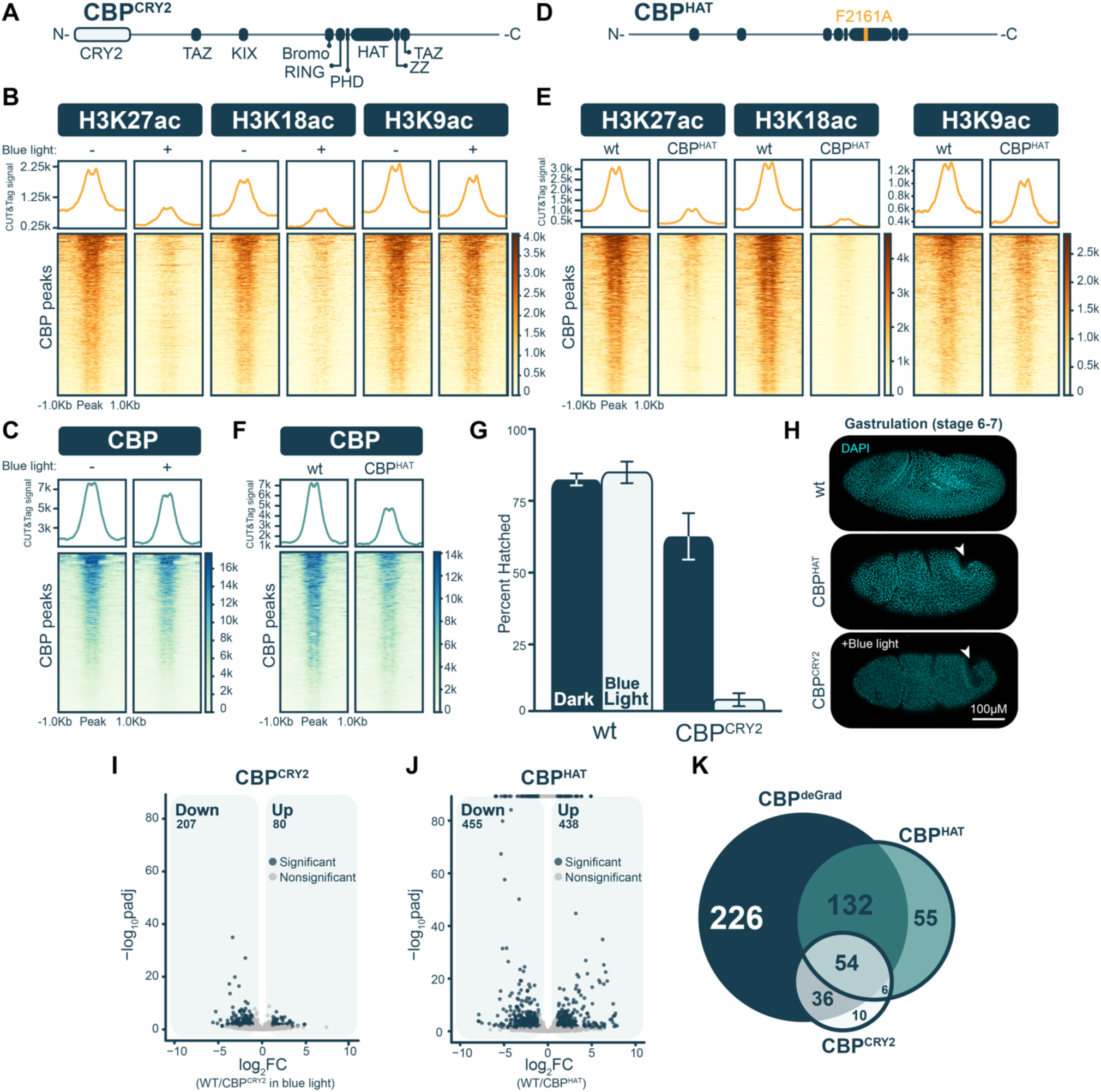
Both catalytic-dependent and independent functions of CBP are required for embryonic development and ZGA. **(A,D)** Diagrams of endogenous CBP^CRY2^ **(A)** and CBP^HAT^ **(D)** with predicted protein domains from UniProt database. **(B,E)** Heatmaps of H3K27ac, H3K18ac, and H3K9ac CUT&Tag from stage 5 CBP^CRY2^ embryos treated in the dark (-) or in blue light (+) **(B)** and wild-type or CBP^HAT^ embryos **(E)** centered on CBP-binding sites. **(C,F)** Heatmaps of CUT&Tag for CBP from stage 5 CBP^CRY2^ embryos treated in the dark (-) or in blue light (+). **(C)** or for CBP from stage 5 wild-type or CBP^HAT^ embryos **(F)**. (G) Hatching rates for wild-type or CBP^CRY2^ embryos raised in the dark (wild type n= 411, CBP^CRY2^ n=429) or treated with blue light (wild type n= 507, CBP^CRY2^ n=400) during the MZT (0-3hrs AEL). Error bars are the standard deviation between replicates. (G) Wild-type, CBP^HAT^, and blue-light treated CBP^CRY2^ embryos at gastrulation stained with 4′,6-diamidino-2-phenylindole (DAPI). Arrowhead points to invagination arrest observed in mutants **(I)** Volcano plot of single-embryo RNA-seq from blue-light treated CBP^CRY2^ embryos as compared to wild-type embryos similarly exposed to blue light. Navy represents genes that change in expression, and grey represents genes with statistically insignificant changes in expression (adjusted p-value < 0.05, |log2 fold change| > 1). **(J)** Volcano plot of bulk RNA-seq from CBP^HAT^ embryos as compared to wild-type embryos. Navy represents genes that change in expression, and grey represents genes with statistically insignificant changes in expression (adjusted p-value < 0.05, |log2 fold change| > 1). **(K)** Venn diagram showing the overlap of decreased, zygotically expressed genes identified in CBP^deGrad^, CBP^CRY2^ with blue light, and CBP^HAT^ embryos.

To determine whether blue-light treatment causes the release of CBP from chromatin and the subsequent decrease in acetylation, we assayed for CBP^CRY2^ occupancy with and without blue-light treatment. Unexpectedly, CBP^CRY2^ occupancy was largely retained after blue-light inactivation (Figure 3C), suggesting blue-light exposure specifically inhibited the catalytic activity of CBP without disrupting its binding. Despite the overall retention of inactivated CBP^CRY2^ on chromatin, a small subset of CBP-binding sites was lost upon blue-light treatment. Our results suggest that blue-light treatment of CBP^CRY2^ results in specific inactivation of the catalytic activity of CBP. To test this further, we generated an allele that specifically inhibits the catalytic activity of endogenous CBP (F2161A within the HAT domain) (Figure 3D)^26,49^. This mutation was not viable as a homozygote, demonstrating that the catalytic activity of CBP is required for development and necessitating the use of germline clones to produce embryos with only the catalytic dead mutant maternally provided (CBP^HAT^). Similar to the CBP^CRY2^ embryos, CBP^HAT^ embryos had reduced levels of H3K27ac and H3K18ac, but not H3K9ac (Figure 3E and S4B). Furthermore, CBP^HAT^ was also largely retained on chromatin with reduced occupancy at a subset of binding sites (Figure 3F). The similarity between CBP^CRY2^ and CBP^HAT^ demonstrates that CBP^CRY2^ allows for optogenetic control of CBP catalytic activity.

We assayed the viability of both CBP^CRY2^ embryos treated with blue light during the MZT and CBP^HAT^ embryos to determine the effects of loss of CBP catalytic activity on early embryonic development. Both conditions resulted in embryos that failed to hatch, and analysis of fixed embryos suggest embryos arrest at gastrulation (Figure 3G,H). It is possible that retention of the catalytic-dead CBP on chromatin dominantly inhibits development, similar to what was reported previously for CRY2-tagged Bicoid^48^. To test this, we assayed hatching rates for progeny from mothers heterozygous for *CBP^CRY2^* whose progeny will inherit both wild-type and CRY2-tagged CBP. There was no difference in hatching rates between blue-light treated and dark controls, suggesting CBP^CRY2^ is not acting as a dominant negative upon blue-light inactivation (Figure S4C). We then used our optogenetic allele to test whether maternally provided CBP activity is required in the early embryo for development. For this purpose, we performed directional crosses in which only one of the parents contributed the *CBP^CRY2^* allele and compared hatching rates of the progeny exposed to either blue light for 0-3 hr AEL or kept in the dark. Progeny from crosses in which the mothers contributed the *CBP^CRY2^* exhibited lethality rates comparable to homozygous *CBP^CRY2^* when treated with blue light (Figure S4D). By contrast, progeny inheriting *CBP^CRY2^* from their fathers, and therefore only possessing zygotically encoded CBP^CRY2^, were unaffected by blue-light treatment (Figure S4E). Thus, maternally encoded CBP catalytic activity is required in the early embryo for progression through the MZT.

### CBP activates gene expression through both catalytic-dependent and independent mechanisms

Knockdown of CBP, either through protein degradation or RNAi, resulted in dramatic changes to the transcriptome during ZGA (Figure 2C)^30^. Nonetheless, it remained unclear whether CBP-mediated gene expression depends on CBP-mediated acetylation. Our *CBP^CRY2^* and *CBP^HAT^* alleles provided powerful tools to determine the catalytic-dependent and independent functions of CBP in genome activation. We performed single-embryo RNA-seq on CBP^CRY2^ embryos treated with blue light from NC10 until 15 minutes into NC14 along with controls (blue-light treated *His2Av* embryos and CBP^CRY2^ embryos kept in the dark). Differential analysis identified 207 genes with decreased expression and 80 with increased expression in CBP^CRY2^ embryos as compared to controls (Figure 3I). As we did previously, we identified genes with changes in expression accounting for effects caused by both the addition of the CRY2 tag to endogenous CBP and treatment with blue light^15^. As in the CBP^deGrad^ embryos, many of the increased genes are maternally expressed (47.7%, 21/42 genes), while the majority of decreased genes are zygotic (66.7%, 88/133 genes) (Figure S4E). These data suggest that CBP-mediated acetylation is important for zygotic gene expression and is supported by bulk RNA-seq from stage 5 CBP^HAT^ embryos (Figure 3J). Compared to CBP^CRY2^, analysis of the bulk RNA-seq on the CBP^HAT^ embryos identified a larger number of genes with changes in gene expression as compared to controls; 438 genes increased in expression and 455 genes decreased (Figure 3J). As before, the increased genes were enriched for those that are maternally expressed while the decreased genes were enriched for zygotically expressed transcripts (Figure 3J and S4F). Differences in the number of differential genes may result from the longer collection period that was sampled in the bulk RNA-seq as compared to the precisely staged single-embryo RNA-seq or the fact that CBP^HAT^ embryos result from germline clones that may affect maternally provided products. Independent of the genotype used to inactivate CBP catalytic activity or the sequencing method utilized, a subset of zygotically expressed genes depend on CBP catalytic activity for transcriptional activation.

While our data demonstrated that CBP catalytic activity is required for activation of a subset of the zygotic genome, we identified fewer genes that changed in expression upon inhibition of CBP catalytic activity as compared to the degradation of the entire protein. When we focused on the zygotically expressed genes, there were 226 genes that were uniquely decreased in the CBP^deGrad^ embryos. Only 16 genes were reduced in the CBP^CRY2^ embryos as compared to the CBP^deGrad^ embryos and 61 in the CBP^HAT^ as compared to the CBP^deGrad^ (Figure 3K). To determine whether the reduction in gene expression in our catalytically inhibited CBP embryos was due to loss of CBP binding at the loci or the absence of acetylation, we determined whether CBP was bound proximally to these genes and, if so, whether binding was maintained in CBP^HAT^ embryos. We then plotted the average log_2_ fold change of these CBP-bound genes. This analysis demonstrated that expression was reduced for genes proximal to CBP-bound regions regardless of whether proximal CBP binding was reduced in the catalytic dead mutants (Figure S4G). Thus, the observed decrease in gene expression cannot be fully explained by the loss of CBP genome occupancy in catalytically dead or inactivated embryos. Together, our analysis of multiple mutants that disrupt CBP function demonstrates that CBP acts through both catalytic-dependent and independent mechanisms to activate expression from the zygotic genome.

### CBP recruits Pol II independent of catalytic activity but catalytic activity is required for release of Pol II from promoter-proximal pausing

To explore how catalytic and non-catalytic CBP activities affect gene expression, we performed spike-in normalized CUT&Tag experiments with antibodies that recognize the initiating form of RNA polymerase II (Pol II) marked by serine 5-phosphorylation on the C-terminal domain of Pol II (Ser5-P). In 2-3 hr old CBP^deGrad^ embryos, the global Ser5-P signal over transcription start sites (TSS) was severely reduced as compared to CBP^GFP^, indicating that Pol II recruitment or stability at the promoter depends on CBP (Figure 4A). Supporting this finding, TATA-binding protein (TBP) was similarly reduced at promoters in 2-3 hr CBP^deGrad^ embryos (Figure 4B). To compare catalytic with non-catalytic functions, we performed similar experiments on 2-3 hr CBP^CRY2^ embryos with and without blue light. In contrast to the results with CBP^deGrad^ embryos, Ser5-P CUT&Tag signal was unaffected over the TSS in CBP^CRY2^ embryos exposed to blue light (Figure 4C). However, Ser2-P, the elongating form of Pol II, was reduced as compared to embryos not exposed to blue light (Figure 4D). These data suggest that catalytically inactive CBP can recruit Pol II to promoters, but this Pol II is not able to release into productive transcription. In early embryos, Pol II is localized to small foci and this localization requires both Zld and CBP^46,50^. By contrast, the more dramatic accumulation of Pol II at the histone locus body is not dependent on these factors. As expected based on these prior studies, staining of control and CBP^deGrad^ embryos showed a similar reduction in Pol II foci upon CBP depletion (Figure 4E). By contrast, Pol II foci were still evident in CBP^CRY2^ embryos exposed and fixed in blue light (Figure 4F). Together our genomic analysis and immunostaining indicate that CBP is required for Pol II recruitment, but that catalytic activity is only necessary for a subsequent step.

**Figure 4.**
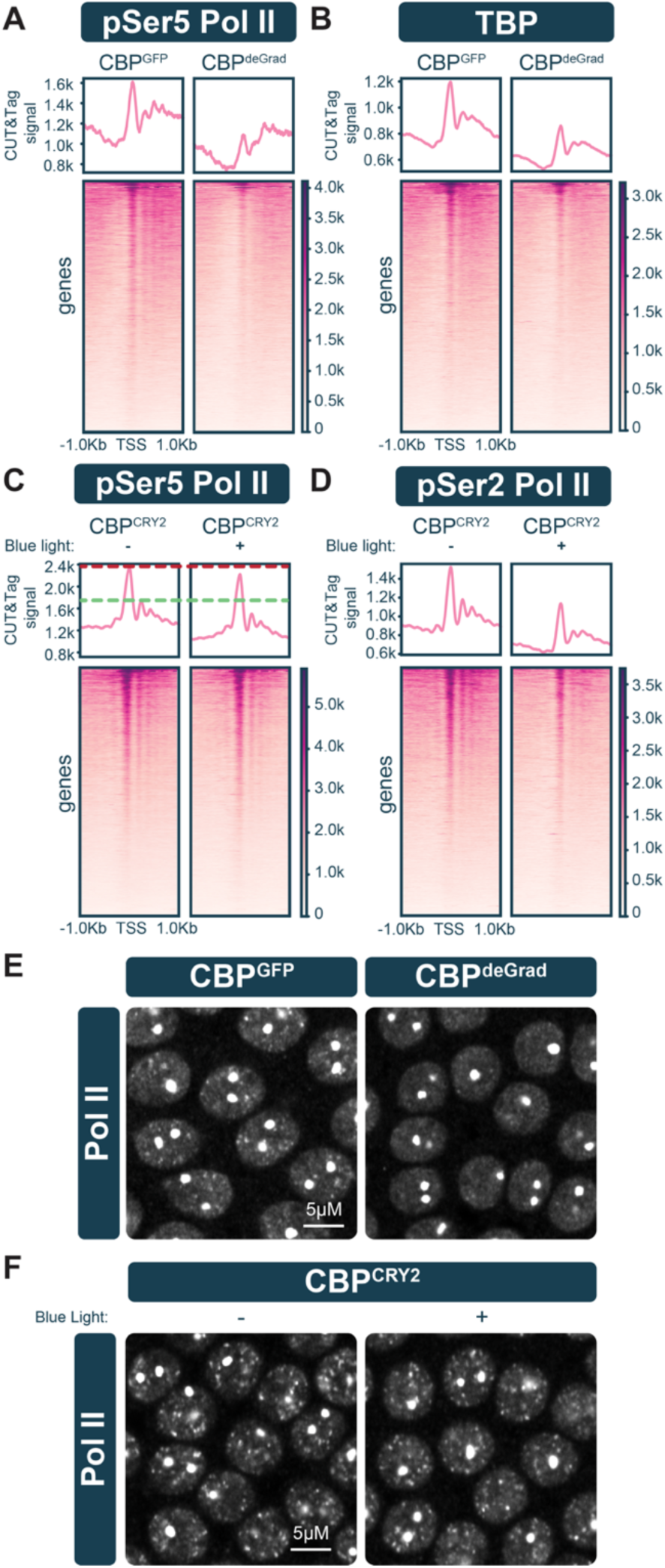
Catalytic activity of CBP is required for RNA Pol II elongation but not recruitment. **(A, B)** Heatmaps of CUT&Tag from stage 5 CBP^GFP^ control and CBP^deGrad^ embryos for pSer5 Pol II **(A)** and TBP **(B)** centered on the TSS of Pol II transcribed genes. **(C, D)** Heatmaps of CUT&Tag from stage 5 CBP^CRY2^ embryos treated in the dark (-) or in blue light (+) for pSer5 **(C)** and pSer2 Pol II **(D)** centered on the TSS of Pol II transcribed genes. **(E)** Staining for RNA Polymerase II in CBP^GFP^ control and CBP^deGrad^ embryos. **(F)** Staining for RNA Polymerase II in CBP^CRY2^ embryos treated in the dark (-) or in blue light (+). Large foci correspond to the histone locus bodies. Scale bars, 5 µm.

To provide a map of transcriptionally engaged Pol II, we performed Precision run-on sequencing (PRO-seq). This analysis identified 207 genes that were transcriptionally downregulated and 54 genes upregulated after spike-in normalization in 2-3 hr CBP^deGrad^ embryos (Figure S5A). While not identical to the gene expression changes identified by mRNA-seq, the gene expression changes identified by PRO-seq correlate well with the single-embryo RNA-seq (Figure S5B). Importantly, PRO-seq captures promoter-proximal paused polymerases. Downregulated genes had Pol II paused at the transcription start site (TSS) in CBP^GFP^ control embryos, whereas unaffected and upregulated genes had little Pol II pausing (Figure 5A, B). Consistent with the Ser5-P and TBP CUT&Tag, Pol II pausing was reduced in CBP^deGrad^ embryos (Figure 5A, B). CBP occupancy was enriched at downregulated gene promoters, but upregulated and unaffected promoters had little CBP. This suggests that downregulated genes are direct CBP targets whereas upregulated genes are indirectly affected by CBP depletion (Figure 5C). These results indicate that CBP is needed to establish paused Pol II by recruiting or stabilizing Pol II at promoters, consistent with earlier findings^51^.

**Figure 5.**
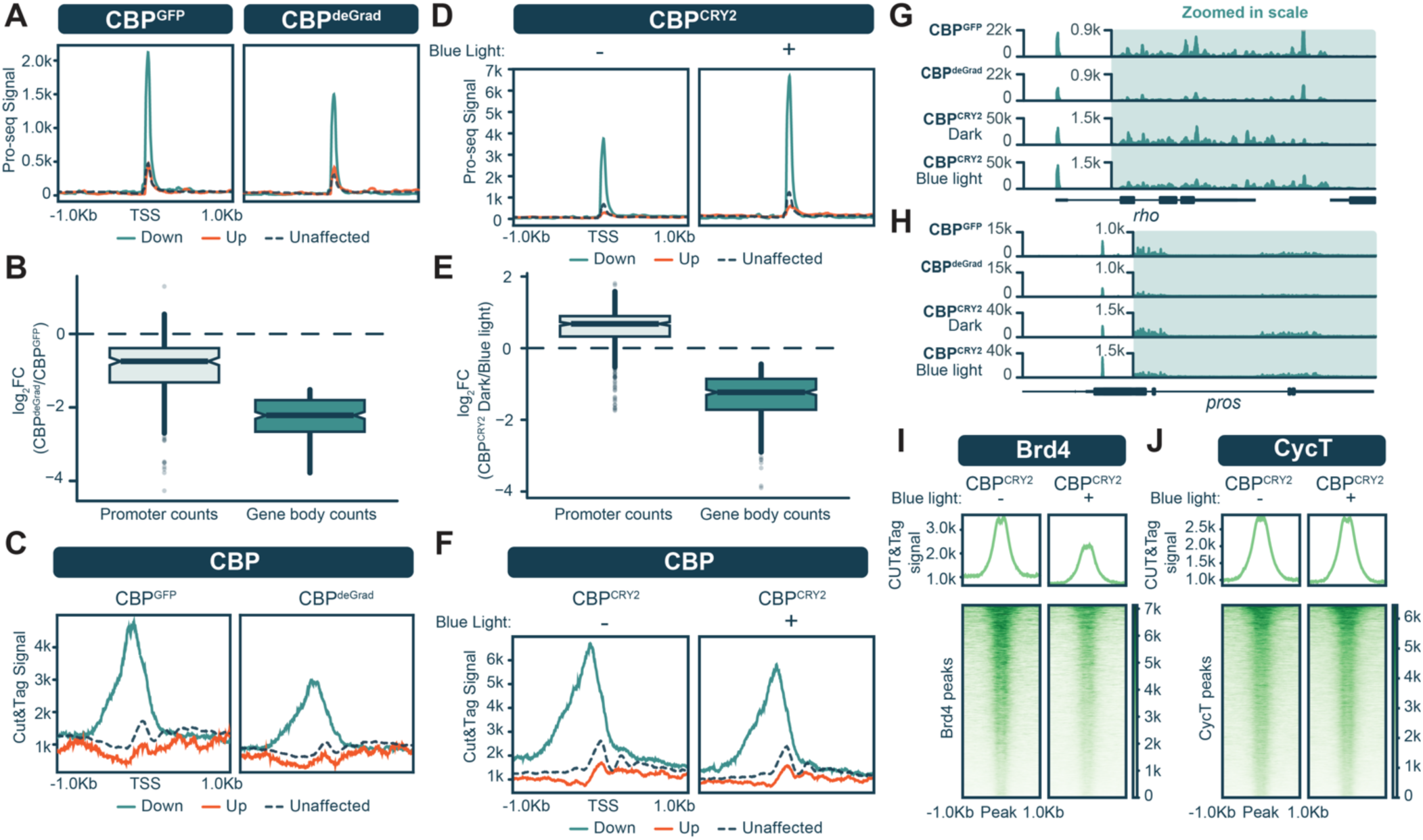
RNA Pol II is retained at promoters in the absence of CBP catalytic activity. **(A)** PRO-seq metaplots centered on the TSS of down-regulated, up-regulated and non-affected genes in stage 5 CBP^GFP^ control and CBP^deGrad^ embryos. **(B)** Box plots of PRO-seq promoter and gene body signal of down-regulated genes in CBP^deGrad^ as compared to CBP^GFP^ controls. **(C)** CBP CUT&Tag signal at promoters of down-regulated, up-regulated and unaffected genes in CBP^GFP^ control embryos. **(D)** PRO-seq metaplots centered on the TSS of down-regulated, up-regulated and unaffected genes in stage 5 CBP^CRY2^ embryos treated in the dark (-) or in blue light (+) (**E**). Box plots of PRO-seq promoter and gene body signal of down-regulated genes in CBP^CRY2^ embryos treated with blue light as compared to untreated CBP^CRY2^ embryos (**F**). CBP CUT&Tag signal in promoters of down-regulated, up-regulated and non-affected genes in CBP^CRY2^ embryos kept in the dark or treated with blue light. (**G,H**). Genome browser tracks showing PRO-seq signal over *rhomboid* (*rho*) **(G)** or *prospero* (*pros*) **(H)** in embryos as indicated to the left. Gene bodies (blue-highlighted regions) are shown with an increased scale (as noted) to enable visualization of PRO-seq reads. **(I,J)** Heatmaps of CUT&Tag for BRD4/fs(1)h **(I)** or CycT **(J)** from stage 5 CBP^CRY2^ embryos treated in the dark (-) or in blue light (+).

PRO-seq on CBP^CRY2^ embryos identified 230 downregulated genes and 315 upregulated genes, which are correlated with changes identified by mRNA-seq (Figure S5C,D). In contrast to the decrease in Pol II pausing observed in the CBP^deGrad^ embryos, promoter-proximal PRO-seq signal increased in CBP^CRY2^ embryos upon blue-light exposure (Figure 5D, E). Despite this increase in Pol II pausing upon blue-light exposure, these CBP-bound genes were down-regulated (Figure 5E, F). This is evident at the *rhomboid* (*rho*) and *prospero* (*pros*) genes where promoter-proximal Pol II is decreased only in the CBP^deGrad^ embryos, but gene body Pol II is decreased in both blue-light treated CBP^CRY2^ and CBP^deGrad^ embryos (Figure 5 G,H). Together these data strongly suggest that Pol II cannot be efficiently released into productive elongation in the absence of CBP catalytic activity. Release of Pol II from pausing depends on P-TEFb, consisting of the Cdk9 kinase and Cyclin T (CycT), and on the bromodomain protein Brd4, also known as female sterile (1) homeotic, fs(1)h, in *Drosophila*^52–55^. CUT&Tag with antibodies recognizing *Drosophila* Brd4 and CycT demonstrated that Brd4 occupancy was reduced, but not absent, in CBP^CRY2^ embryos (Figure 5I). Indeed, CycT remained bound to promoters in CBP^CRY2^ embryos despite the reduction in Brd4, suggesting that P-TEFb is recruited in an inactive state independent of Brd4 (Figure 5J). These results support a model that identifies two distinct activities for CBP in zygotic genome activation: CBP establishes paused Pol II by a catalytically independent mechanism, whereas catalytic activity is important for release of paused Pol II into elongation downstream of P-TEFb recruitment.

### The pioneering activity of Zld does not depend on CBP

Our data identify two functions for CBP in promoting gene expression but do not define the relationship between CBP and pioneer activity. Because we showed that Zld recruits CBP to activate zygotic gene expression, we wanted to test if CBP-mediated acetylation was required for Zld to promote chromatin accessibility. To test this, we induced Zld expression in S2 cells and treated with either A-485, a specific catalytic inhibitor of CBP, or DMSO, as a control, and performed ATAC-seq^56^. We confirmed catalytic inactivation of CBP using ChIP-seq and immunoblots to assay for H3K27ac levels, which were globally decreased at all Zld-bound sites (Figure S6A,B). In contrast to the dramatic reduction in acetylation, chromatin accessibility was unchanged at Zld-pioneered regions (Figure 6A), indicating that histone acetylation is not necessary for Zld-mediated accessibility at these sites. We then tested the role of acetylation in embryos by performing single embryo ATAC-seq on CBP^CRY2^ embryos exposed to blue light and compared chromatin accessibility at CBP-bound sites to *His2AvRFP* controls also exposed to blue light. Similar to our results in culture, we did not identify global changes in accessibility at either Zld-bound or CBP-unique loci (Figure 6B). Quantitative analysis identified 5,757 sites with changes in accessibility between CBP^CRY2^ and control embryos (1454 increased, 4303 decreased), but only 7.88% were bound by CBP, suggesting these are indirect effects (Figure S6C). Our analysis in cell culture and embryos demonstrates that CBP-mediated histone acetylation is not required for Zld pioneer activity.

**Figure 6.**
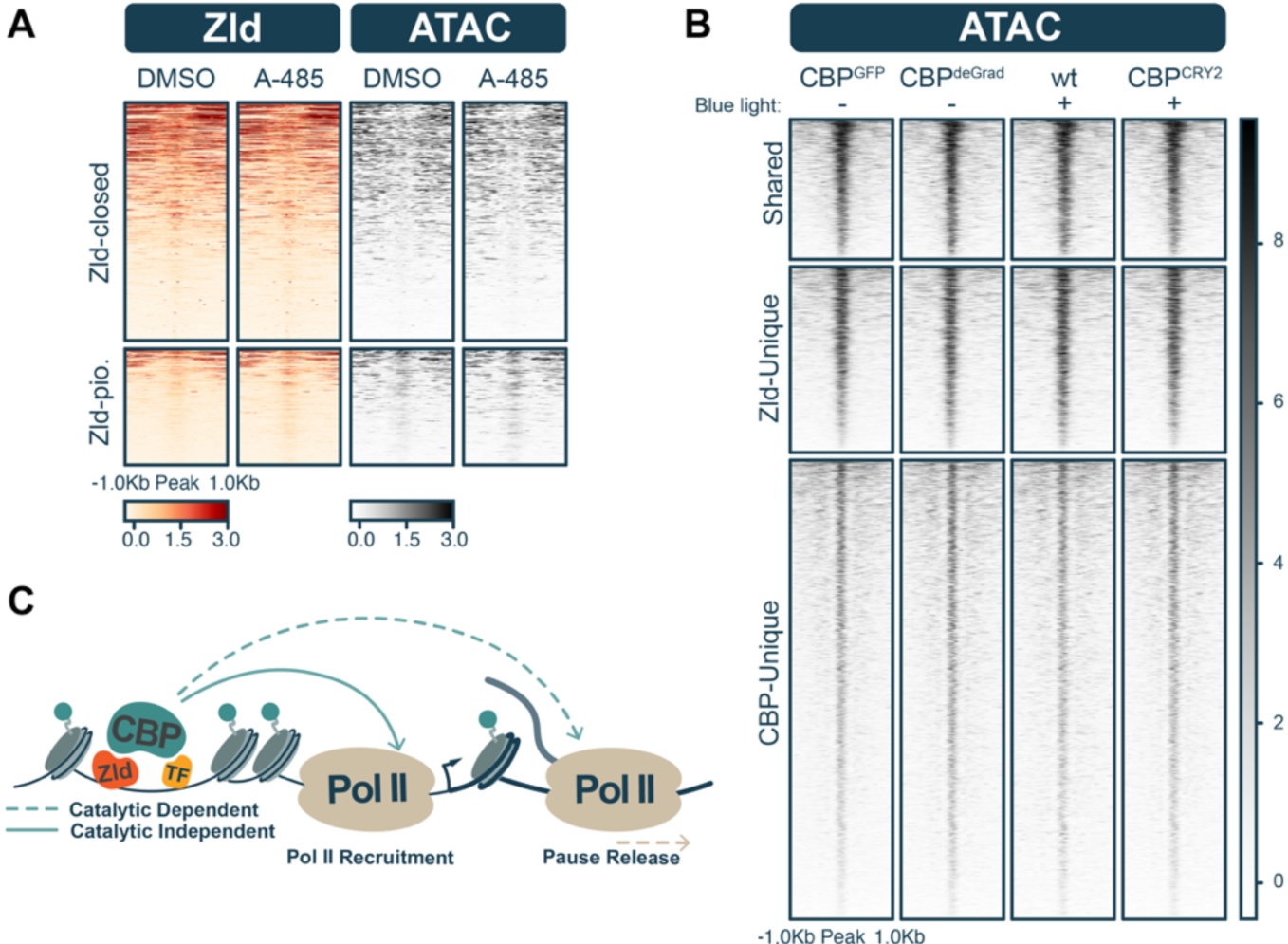
CBP is not required for Zld-mediated pioneer activity. **(A)** Heatmaps of Zld occupancy (ChIP-seq), chromatin accessibility (ATAC-seq) in S2 cells treated with either DMSO or A-485 divided into regions that Zld pioneers (Zld-pio.) and those that remain inaccessible (Zld-closed). Signals are z-score normalized. **(B)** Heatmaps of single embryo ATAC-seq with peaks sorted into shared, Zld-unique, and CBP-unique classes in wild-type, CBP^deGrad^ or CBP^CRY2^ embryos (as labeled). Embryos were staged precisely 15 min into NC14 before harvesting. Plus and minus signs denote whether embryos were treated with blue light from NC10-14. Signals are z-score normalized. **(C)** Model depicting Zld recruitment of CBP to *cis-*regulatory regions in the early embryo. Independent from its role as a histone acetyltransferase, CBP is needed to recruit Pol II. By contrast, CBP catalytic activity is required for Pol II elongation.

To investigate if CBP is more generally important for chromatin accessibility, we leveraged our deGrad system to perform *in vivo*, single-embryo ATAC-seq on CBP^GFP^ and CBP^deGrad^ embryos precisely 15 min into NC14. Similar to what we observed when CBP catalytic activity was inactivated but CBP occupancy was retained, chromatin accessibility was largely unchanged at Zld-bound and CBP-unique sites (Figure 6B). Quantitatively calling differential sites identified only 1,094 sites (671 up, 423 down) that changed in accessibility in CBP^deGrad^ embryos as compared to controls and only 8.87% (97 sites) overlapped with CBP^GFP^ ChIP-seq (Figure S6D). Thus, CBP and CBP-mediated acetylation are not broadly required for chromatin accessibility during the MZT.

## Discussion

We demonstrated that CBP is required for zygotic transcription during the MZT, and its recruitment is dependent upon the pioneer factor Zld at shared *cis*-regulatory sites. Two mechanisms could explain this Zld-mediated recruitment of CBP. Zld might indirectly recruit CBP through its pioneer activity by promoting the binding of an additional transcription factor that directly interacts with CBP. Possible candidate transcription factors are Bcd and Dl, which directly interact with CBP and depend on Zld for recruitment^12,13,40,57–59^. Our motif analysis supports this model, as we identified Bcd motifs enriched at Zld/CBP shared regions and Dl motifs enriched more broadly at CBP-bound sites. Alternatively, Zld might directly recruit CBP to shared sites either through interactions with CBP or with an intermediate cofactor. Direct interactions between Zld and CBP have not been reported. Nonetheless, Zld is sufficient to induce H3K27ac at Zld-pioneered regions in S2 cells, where Bcd and Dl are not expressed. Thus, in S2 cells Zld might directly promote CBP binding or could function through another transcription factor that is endogenously expressed in S2 cells. Further experiments will be required to distinguish between these two models.

Our identification of CBP-bound regions that were distinct from Zld-bound regions suggests that CBP recruitment is not solely dependent upon Zld. Those sites uniquely bound by CBP were enriched for motifs bound by the insulator binding proteins Dref and BEAF-32, suggesting that CBP might play an important role in establishing tertiary chromatin structures. Indeed, CBP is known to bind outside of active cis-regulatory regions, including insulators where CBP-mediated acetylation of H3K27 blocks the spreading of silencing H3K27me3 into euchromatic regions^38^. Catalytic inhibition of CBP/p300 in cell culture disrupts enhancer and promoter interactions^60^. Uniquely bound CBP sites may therefore play a distinct role in maintaining higher order organization of the genome.

Degradation of maternally encoded CBP results in the downregulation of over a thousand genes during widespread genome activation. Our finding is supported by orthogonal approaches of CBP mediated knock down at the transcript level rather than protein level^30^. Many of the genes that depend on CBP are also bound and dependent upon Zld for activation, indicating CBP works in conjunction with Zld to upregulate transcription of the zygotic genome. Coordinated activation of these targets is likely a stepwise process, where Zld first binds to the genome and recruits CBP, which stimulates the downstream recruitment of RNA Pol II to promoters. Formation of transcriptional hubs has been suggested to increase local concentrations of transcriptional regulators at promoters to drive transcription^61^. High-resolution microscopy resolved the formation of Zld-dependent transcriptional hubs^62,63^, and removal of Zld from the nucleus or knock down of CBP by RNAi abolishes RNA Pol II cluster formation^46,50^. Our work provides mechanistic details, demonstrating that CBP and Zld can bind to the same genomic loci, that CBP binding depends on Zld, and that both factors are required for transcriptional activation.

The CBP/p300 HAT family is multifaceted in its coactivator function where CBP and p300 can activate transcription through acetylation of histone tails, acetylation of transcriptional coactivators, or independent of HAT activity by acting as a transcriptional bridge between enhancers and promoters^64^. We showed that CBP mediates gene expression through catalytic-dependent and independent mechanisms (Figure 6C). A majority of acetylation-dependent genes overlapped with genes that require CBP for activation, but not vice versa. Using RNA-seq, we identified a more dramatic reduction in transcription in embryos with a reduction in CBP protein as compared to those in which only catalytic activity was disrupted. These data indicate that CBP-mediated transcriptional activation is largely dependent on functions separate from catalytic activity. Possible mechanisms of catalytic-independent activation by CBP include the recruitment of general transcription factors, like TFIIB, to help the formation or stabilization of the pre-initiation complex of RNA Pol II^51,65–67^. This is consistent with the reduction in Ser5-P Pol II and TBP at promoters and lack of Pol II foci in CBP-depleted embryos. Alternatively, CBP-mediated histone acetylation has been shown to facilitate recruitment of bromodomain containing coactivors such as Brd4 or TBP^46,68^. Thus, CBP facilitates the recruitment of the transcription machinery through multiple diverse functions. This is similar to what has recently been shown for other histone modifiers and highlights the diversity of functions that can be carried out by these large enzymes^69^.

Despite the milder effects on gene expression of catalytic inactivation as compared to CBP depletion, our data demonstrate that the catalytic function of maternally encoded CBP is essential for embryogenesis; embryos are nonviable when we deplete maternal CBP protein or inactivate catalytic activity. This contradicts a recent report in which transgenic overexpression of the F2161A catalytic dead allele of CBP restored viability to embryos in which CBP was depleted through RNAi^30^. Immunoblots for CBP showed considerable loss of CBP protein after RNAi knock down, however, there were detectable levels of CBP. We propose that low levels of wild-type catalytically active CBP protein present in these knockdown embryos was sufficient to restore viability with the presence of ectopically expressed catalytic dead CBP. The reproducibility of results observed in both CBP^CRY2^ inactivated and CBP^HAT^ mutant embryos strongly supports our conclusion that catalytic activity of maternally provided CBP is essential during ZGA.

We find that CBP-target genes contain promoter-proximal paused Pol II and that CBP catalytic activity is needed for release of paused Pol II into productive elongation. In agreement with this, pause release can be stimulated by increased acetylation^70^. In embryos without CBP catalytic activity paused genes become more strongly paused, loose Ser2-P Pol II, and nascent transcription over gene bodies is decreased. Loss of acetylation results in impaired binding of BRD4, but not P-TEFb. Although phosphorylation of Pol II and negative elongation factors by P-TEFb is necessary for release into elongation^42^, BRD4 is also needed for Pol II pause release independent of P-TEFb recruitment^71–73^. Since BRD4 recognizes acetylated histones^25^, a plausible scenario is that CBP-mediated histone acetylation allows BRD4 to engage with paused zygotic genes to stimulate pause release.

Zld was the first pioneer factor identified as a master regulator of zygotic transcription across metazoans and has provided a paradigm in understanding how the genome is reprogrammed during early embryo development. Nonetheless, the mechanisms by which Zld mediates the displacement of nucleosomes and activates zygotic transcription remain unclear. Prior to this study, we hypothesized that recruitment of HAT cofactors by Zld might promote chromatin accessibility as histone acetylation has been correlated with active chromatin^33,74^. In contrast to this expectation, our data clearly separates CBP-mediated gene expression from chromatin accessibility and demonstrate that the ability of Zld to open chromatin is independent from CBP recruitment, consistent with earlier studies on the relationship between CBP and chromatin accessibility^75^. Screening for potential Zld-recruited cofactors may help shed light on how Zld mediates chromatin opening. However, it is possible that Zld promotes chromatin accessibility through creating a transient hub that concentrates both itself and other factors and in so doing promotes chromatin occupancy. Our work makes it evident that pioneering function and transcriptional activation are separable and that to uncover the mechanism of Zld pioneer function it will be essential to focus on chromatin accessibility apart from gene expression.

While Zld has served as a paradigm for understanding pioneer-factor regulated genome activation, Zld is not conserved outside of the Pancrustacea clade of arthropods^76^. By contrast, CBP is broadly conserved^77,78^. Similar to our demonstration that Zld recruits CBP to activate gene expression, pioneer factor mediated recruitment of p300/CBP initiates ZGA in vertebrates. In zebrafish, Nanog, Pou5f3, and Sox19b activate zygotic transcription and recruit p300a, which is essential for ZGA^21,22,79^. As in flies, histone acetylation is separable from chromatin accessibility^22^. Members of the Dux transcription factor family initiate ZGA in mammals^80,81^. Dux4 in humans directly interacts with p300, where removal of the Dux4-p300 interacting domain results in a failure to activate transcription^82^. Furthermore, inhibition of p300/CBP catalytic activity in mouse embryos impedes zygotic transcription^83^. Thus, while the individual pioneer factors are not conserved across model systems, they share a conserved role in recruiting CBP/p300 to activate zygotic transcription. Our work describes a model by which this conserved HAT coordinates the activation of the zygotic genome in tandem with a pioneer factor to jump start development and identifies roles in both Pol II recruitment and release.

## Acknowledgements

We would like to thank Renato Paro, Nicola Iovino, Kazuko Hanyu Nakamura, and Jim Kadonaga for sharing antibodies used in the study. We also thank the Bloomington Drosophila Stock Center and the Drosophila Genome Resource Center for providing reagents and fly lines. We acknowledge the University of Wisconsin-Madison Biochemistry Department Optical Core and Imaging facility at Stockholm University (IFSU) for access to microscopes and the University of Wisconsin-Madison Biotechnology Center, the NUSeq Core Facility, Bioinformatics and Expression Analysis (BEA) core facility at Karolinska Institutet, and SciLifeLab, Stockholm for sequencing. AJM was supported by an NSF graduate research fellowship. Experiments were supported by a NIH R35 GM136298 (MMH), Swedish Research Council, 2022-03650 and Cancerfonden, 23 2959 Pj grants (MM). MMH was also supported by a Vallee Scholar Award. MMH is a Romnes Faculty Fellow and Vilas Faculty Mid-Career Investigator.

## Author contributions

AJM, SP, TJG, GH, MMH, and MM designed the experiments. AJM, SP, AJR, SS, TJG, GH, YK performed the experiments and data analysis. AJM, SP, MMH, and MM wrote the original draft. AJM, SP, TJG, MMH and MM revised and edited the manuscript. AJM, MMH, and MM acquired funding.

## Data availability

All sequencing data is available through GEO accession number GSE276949.

## Materials and Methods

### Drosophila husbandry and strains

All stocks used in this study were maintained at 25°C and fed a molasses or potato mash diet. Non-CRY2 lines were raised on a 12-hour light and dark cycle. CBP^CRY2^ flies were laid and hatched in the dark. Stocks are listed in the Reagents Table.

### Cell culture

The stable S2 MT-Zld line used in this study was described previously^16^. Cells were cultured in Schneider’s medium (Thermo Fisher Scientific) supplemented with 10% FBS (Omega Scientific) and 10% antibiotic-antimycotic (Thermo Fisher Scientific). Puromycin (Fisher Scientific) was added to media to a final concentration of 1 µg/ml to select for transgenic cells.

### Cas9-mediated genome engineering

The N-terminus of CBP was endogenously tagged with GFP or CRY2 using a previously described method for Cas9 mediated genome engineering^84^. The double stranded donor used as a template for homologous recombination was generated via Gibson assembly (NEB). Sequence encoding either GFP or CRY2 was added immediately after the start codon of the *CBP* open reading frame and flanked by 1kb homology arms. For screening purposes, a *3XP3-DsRed* cassette was inserted into the first *CBP* intron and flanked by long terminal repeats recognized by the PiggyBac transposase. The guide RNA sequence (GTCAAACTAACTCTATATGA) was cloned via inverse PCR into pBSK downstream of a U6 promoter. Donor and guide RNA plasmids were purified and sent to The Best Gene Inc. for injection into *w[1118]; PBac{y[+mDint2]=vas-Cas9}VK00027* embryos. Lines were then screened for DsRed positive eye expression to identify Cas9-edited flies, followed by excision of the DsRed cassette by the PiggyBac transposase. To validate proper editing, each founder line was confirmed by sequencing.

A point mutation that disrupts CBP catalytic activity^49^, F2161A, was generated by a two-step CRISPR- and recombination mediated cassette exchange (RMCE)-based strategy^85^. Two gRNAs and a 3xP3-DsRed containing donor plasmid with attP sites were injected into *vas-Cas9* flies to delete exons 9 and 10 and the intervening intron (1.4 kb). To select the gRNAs, *vas-Cas9* genomic DNA from flanking introns was amplified with primers 5pSequenceFw, 5pSequenceRw, 3pSequenceFw and 3pSequenceRw and sequence verified. The gRNAs were cloned into pCFD3. Around 1 kb long homology arms were amplified with primers CBP HAL-Fw, CBP HAL-Rw, CBP HAR-Fw and CBP HAR-Rw and subcloned in the pJET 1.2 vector by CloneJET PCR blunt end cloning (Thermofisher Scientific K1231). The resulting plasmid was combined with plasmid pJET1.2-STOP-dsRed by Golden Gate assembly using *BsmBI* to make the final pBS-STOP-dsRed dCBP donor plasmid. The pCFD3 gRNAs (110ng/ul each) and pBS-STOP-dsRed dCBP plasmid (500ng/ul) were injected into *w[1118]; PBac{y[+mDint2]=vas-Cas9}VK00027* embryos. Four females with dsRed positive eye expression were recovered and two of these had ends-out targeting of the donor plasmid as determined by PCR. A stock from one of these *CBP^RMCE^* flies was injected with a vasa-phiC31 plasmid and attB-flanked 1.4 kb genomic DNA containing exons 9 and 10 with the F2161A substitution. For the attB-dCBP plasmid, primers AmpCBP2step-Fw and AmpCBP2step-Rw were used to amplify genomic DNA and cloned at *HindIII* and *KpnI* restriction sites of pABC 2.0. The desired nucleotide changes (TTT to GCC, F2161A) were introduced by site directed mutagenesis with primers SiteD-CBPFw and SiteD-CBPRw. F1 progeny were screened for absence of dsRed, and correct orientation of the genomic DNA after RMCE confirmed by PCR. This *CBP^HAT^* stock was balanced with FM7h. To rule out splicing defects by the inserted intronic attR sequences, cDNA was generated and primers CBPForward and CBPRev1 used to amplify the exons which were sequence verified. To ensure that no other lethal mutations were present in the stock, it was crossed to a heat shock CBP transgenic line, and heat shock-induced CBP expression was able to rescue viability.

To introduce a FRT sites into this line, we mated *CBP^HAT^*/FM7h flies with FRT strain *P{w[+mC]=Ubi-mRFP.nls}, w1118, P{ry[+t7.2]=neoFRT}19A* (Bloomington #31416) for meiotic recombination. We confirmed successful recombination by PCR detection of the FRT insertion and Sanger sequencing of the F2161A mutation. To obtain homozygous CBP^HAT^ mutant embryos we used the dominant female sterile technique. We crossed *CBP^HAT^ FRT*/FM7h females with *ovoD1 hsFLP FRT* (Bloomington #23880) males. Progeny from this cross were heat-shocked at 37°C for 2 h 15 min on days 3, 4, and 5 after egglaying. Embryos were collected on apple juice plates supplemented with fresh yeast and aged at 25°C for specific time ranges dependent on the particular experiment, detailed in the relevant methods section.

### Hatching rates

Embryos were collected on molasses agar plates fitted to embryo collection cages (Genesee Scientific). Following a one-hour pre-lay, the plate was replaced and embryos were collected. 1 hr after egg laying (AEL) embryos were picked from the plate and aligned in the middle of a new plate. More than 24 hours AEL, unhatched embryos were counted and used to calculate the percent embryos viable from each experiment. At least three replicates were performed for each experimental genotype. CRY2 control experiments were laid and aged in the dark. Blue-light treated CBP^CRY2^ embryo cages were set up in a cardboard box fastened with blue LED light strips for an hour lay. Following the hour, the molasses collection plate was removed and left in the blue-light box for another 2 hours to ensure blue-light inactivation through the MZT. After a total of 3 hours, embryos were aligned on a new plate and left to age for the remainder of the 24 hours in the dark.

### Western blots

Protein lysates were transferred from a polyacrylamide gel (6% for Zld, 8% for tubulin, 15% for histones) to a 0.45µm pore PVDF membrane (Fisher Scientific) for Zld or tubulin, and a 0.2µm nitrocellulose membrane (Pall Life Sciences) for H3K27ac immunoblotting. Transfers to PVDF membranes was done in a 20% methanol, 25mM Tris, 200mM glycine buffer for 75min at 500 mA. Transfer buffer for nitrocellulose membranes was supplemented with 0.375% SDS and transferred for 2 hours at 280 mA. Transfer of CBP was done overnight in 10% methanol, 25 mM Tris, 200 mM glycine buffer with 0.05% Tween 20 at 25V. Membranes were blocked for 30min at room temperature, then incubated overnight with primary Zld^6^ (1:750), tubulin (Sigma, 1:5000), H3K27ac (Active motif, 1:1000), CBP^58^ (1:250), H3K18ac (Abcam, 1:750), H4 (Abcam 1:1500), or H3 (Abcam 1:1000) antibody at 4°C. Secondary antibodies goat anti-rabbit IgG-HRP (BioRad) and goat anti-mouse IgG-HRP (BioRad) were used at 1:3000 and incubated at 1 hour at room temperature. Activation of the chemiluminescent reaction was stimulated using the SuperSignal West Pico Plus kit (Fisher Scientific) and imaged on film or digitally on the Azure Biosytems 600 imaging system. Fluorescent secondary antibodies used were goat anti-rabbit IRDye 680RD (LI-COR, 1:10000) and goat anti-mouse IRDye 800CW (LI-COR, 1:5000) and were imaged on the Odyssey FC (LI-COR Biosciences) dual-mode imaging system.

### Immunostaining and confocal microscopy

For DAPI staining, embryos were collected on apple juice agar plates supplemented with yeast for two hours and aged for two hour (2-4 hr AEL). The embryos were washed, dechorionated in bleach, and fixed in formaldehyde as previously described^86^. DAPI was used at a concentration of 1 μg/mL. Images were acquired with a Zeiss LSM 780 confocal microscope using a 20x objective. Staining for Pol II was executed as described before^50^. Embryos were dechorionated with bleach, fixed in 4% formaldehyde mixed with heptane for 25 min, blocked for 30 min in 1x normal goat serum, and incubated overnight in anti-RNA Pol II-647 conjugate antibody (Sigma, 1:100). The next day, primary stained embryos were washed 4x with 0.1% PBS with Triton-X100 for 30 mins, and stained with DAPI for 30 mins in 50% glycerol. Finally, embryos were transferred to 75% glycerol and mounted onto a slide for imaging on a Nikon A1R+ confocal. Pol II staining images were taken using a 100x objective at a 4.48 zoom. The 647 images were taken using a laser power of 1 and a line averaging of 2.

*CBP^GFP^;His2av-RFP and CBP^deGrad^; His2av-RFP* embryos were live imaged on a Nikon A1R+ confocal provided through the University of Wisconsin-Madison Biochemistry Optical Core. Prior to imaging, 2-3hr embryos were dechorionated in 50% bleach for 3min then mounted on a hydrophobic membrane with halocarbon oil. Imaging was done on a 60x objective in which a single plane was used to capture GFP and RFP fluorescence via 488 and 560 lasers, respectively. Processing of images was done in FIJI^87^.

### ChIP sequencing

The ChIP protocol used on fixed embryos in this study was adapted from a published protocol^88^. Embryos were collected 2-3hrs AEL, dechorionated in 50% bleach for 3 min, and followed by a 15min fixation in formaldehyde (.45% formaldehyde for Zld and H3K27ac, 1.8% for CBP^GFP^). Two replicates of 1000 stage five embryos per experiment were hand sorted under a dissecting scope to avoid contamination by older embryos. Sorted embryos were homogenized in 1mL of RIPA buffer (50 mM Tris-HCl pH 8.0, 0.1% SDS, 1% Triton X-100, 0.5% sodium deoxycholate, and 150 mM NaCl) and underwent 11 cycles of sonication for 20s at 20% output on full duty cycle. For CBP^GFP^ ChIP, a spike-in of fixed and sonicated H3.3-GFP MEF chromatin was added before removing 5% input. IPs were incubated with 6µg antibody overnight at 4°C and purified using Protein A magnetic beads (Invitrogen). After washing and eluting purified IP, DNA from IPs and inputs were treated with 90µg of RNAseA for 30mins at 37°C and decrosslinked by adding 100µg of Proteinase K overnight at 65°C. DNA was purified using a phenol:chloroform extraction with an overnight ethanol precipitation step to concentrate samples. Libraries for inputs and IPs were prepared using the commercial NEB Next Ultra II kit.

ChIP in S2 cells was performed as previously described^16^. 5x10^7^ cells were fixed in .8% formaldehyde for 7 min. Fixed chromatin was sonicated on a Covaris S220 for five 120s rounds with 60s delay, peak power at 170, a duty factor of 10%, and 200 cycles per burst. A spike-in of 5% MEF chromatin was added to H3K27ac sonicated chromatin before separating IPs from 5% input. IPs were incubated with antibody at 4°C for 4 hours. IP purification, RNAse treatment, decrosslinking, and DNA concentration was performed identically to the embryo ChIP protocol stated above.

All ChIP samples were prepared using the NEB Next Ultra II kit (NEB). Samples were sent to the NUseq Core Facility at Northwestern and sequenced on the Illumina HiSeq or Illumina NextSeq 500 using single end 50bp or 75bp reads, respectively.

### ChIP-sequencing analysis

Reads were aligned to the *Drosophila melanogaster* genome (dm6) using bowtie2 v2.4.4^89^. Reads mapped to mitochondrial or scaffold DNA, aligned multiple times, or were unmapped were discarded. MACS2 v2.2 was used to call peaks between each IP and its corresponding input^90^. The BEDtools intersectBed function was used to define high confidence peaks present in both replicates^91^. All bigwigs were generated using bamCoverage from deepTools v3.5.1^92^ and were z-score normalized by taking the average read count across all bins genome wide and dividing by the standard deviation. IPs for H3K27ac in S2 cells and CBP^GFP^ were spike-in normalized in which a scaling factor was determined by the ratio of percent MEF chromatin aligned in the IPs as compared to input.

### Imaging of live embryos for single embryo sequencing

Collection and processing of single embryo replicates for RNA-seq or ATAC-seq was performed as described previously^15,93^. Embryos were collected and washed prior to mounting in halocarbon oil on a glass bottom dish. Real time staging of embryos was possible through the His2av-RFP marker used to measure nuclear density (nuclei/2500µm^2^). Live imaging was performed on a Nikon Eclipse Ti2 inverted fluorescent microscope. Embryos were processed for ATACseq or RNA-seq exactly 15 min into NC14. For optogenetic experiments, His2Av-RFP and CBP^CRY2^;His2Av-RFP embryos were treated with a repeating sequence of 60s exposure to 470nm filtered light followed immediately by a 10 ms exposure to 555nm. This sequence was initiated upon entry into NC10 and ended 15 min into nuclear-cycle 14. The single embryo was transferred from the imaging disc into 40µL Trizol and pierced with a sterile needle to free RNA into solution. Individual samples were then frozen and stored at -80°C until all replicates were collected and ready for RNA extraction. Embryos for ATAC-seq were immediately processed for tagmentation upon reaching 15 mins into NC14.

### Bulk mRNA sequencing

Bulk embryo collections used for the CBP^HAT^ sequencing were collected at 2h15min -3hr AEL. Three replicates were performed between CBP^HAT^ and wild-type controls. After collection, embryos were dechorionated in bleach and added to Trizol. After chloroform extraction, RNA was purified with RNEasy Mini Elute Clean up Kit (Qiegen). Libraries were prepared with the TrueSeq Stranded mRNA kit and sequenced on a NovaSeq 6000.

### Single-embryo mRNA sequencing

Embryos frozen in Trizol at -80°C were homogenized with a sterile pestle. After, 960µL of Trizol supplemented with 200µg/mL glycogen was added. RNA was extracted, cleaned, and concentrated through ethanol precipitation. Libraries were prepared using the Illumina TrueSeq RNA prep kit V2. And sequenced either on the Illumina HiSeq 4000 using single-end 50bp reads or the NovaSeq X Plus using paired-end 150bp reads.

### RNA-sequencing analysis

Alignment of reads to the *Drosophila melanogaster* dm6 reference genome was done with HISAT2 v2.1.0 with a mapping quality threshold of ≥ 30^94^. Reads below this threshold were rejected as were reads that mapped to the mitochondrial or scaffold DNA. Aligned reads were annotated to genes using FeatureCounts 2.0 from Subread^95^. Differential analysis between RNA-seq samples was performed using DEseq2 with statistically significance was defined at padj, 0.05 and |log_2_ fold change| > 1^96^. Changes in gene expression after blue-light treatment that were not specific to CRY2-mediated inactivation were subtracted out by comparing the log_2_ fold change of CBP^CRY2^ embryos to blue-light treated His2Av-RFP control embryos. Changes in gene expression due to the addition of the CRY2 tag alone to the CBP protein were removed by identifying expression changes in CBP^CRY2^ dark embryos over His2av-RFP dark controls. Methods of subtracting changes in gene expression not specific to blue-light inactivation has been described at length previously^15^. For CBP^HAT^ the analysis was performed using the nf-core/rnaseq best-practice analysis pipeline (10.5281/zenodo.1400710).

### ATAC sequencing

Staged single embryos were picked and homogenized in 10µL of pre-chilled lysis buffer (10mM Tris pH 7.5, 10mM NaCl, 3mM MgCl_2_, 0.1%NP-40). Embryo homogenate was resuspended by adding an additional 40µL of lysis buffer and spun down at 500g for 10 min at 4°C. The supernatant was removed under a dissecting scope to ensure the small pellet was not disturbed. 2.5µL of water, 2.5µL of Tn5 enzyme, and 5µL of TD buffer (Illumina Tagment DNA Enzyme and Buffer Kit) were added to resuspend the pellet, followed by a 37°C incubation for 30 min. Libraries were prepared using IDT for Illumina Nextera DNA Unique Dual indices, and PCR amplified with NEBNext Hi-Fi 2x PCR Master Mix (72°C for 5min, 98°C for 30s, followed by 12 cycles of 98°C for 10s, 63°C for 30s, and 72°C for 1min). Libraries were DNA purified using magnetic axygen beads and washed with ethanol (Thermo Fisher Scientific).

ATAC-seq in S2 cells was carried out similarly, in which 200,000 cells were spun down for 3min at 600g at 4°C, washed with 1X PBS, and resuspended in 100μL lysis buffer. Lysed cell material was pelleted after centrifuging for 10min at 600g at 4°C. The supernatant was replaced with 47.5UL TD buffer and 2.5 Tn5 enzyme (Illumina) and incubated at 37°C for exactly 30min. Libraries were prepared exactly as described above.

Sequencing of these ATAC libraries was done at the University of Wisconsin Madison Biotech Center on the NovaSeq6000 with paired-end 150bp reads.

### ATAC-sequencing analysis

ATAC analysis on the other datasets was performed as described previously^16^. Briefly, Adapter sequences were trimmed from reads using NGmerge v0.3^97^. Bowtie2 v2.4.4^89^ was used to align trimmed reads to the dm6 reference genome with a mapping quality threshold of ≥ 30. Reads with a poor mapping quality or aligned to mitochondrial or scaffold DNA were thrown out. For analysis, only fragment sizes smaller than 100bp were used. Reads were merged between replicates before using MACS2 v2.2^90^ to call peaks. Bigwigs were zscore normalized between experiments and differential analysis was performed using DEseq2^96^ with the same parameters described in the RNA-seq analysis description.

### CUT&RUN

CUT&RUN was used to identify H3K27ac deposition in S2 cells using the EpiCypher CUTANA kit and protocol. 2x10^5^ cells were extracted and treated with 0.5μL of H3K27ac antibody or an IgG control overnight at 4°C. After immunopurification, libraries were generated using the NEBNext Ultra II library kit (NEB) and sequenced on an Illumina NovaSeq 6000 with150bp paired-end reads.

### CUT&RUN analysis

Reads were trimmed and aligned as described in the ATAC-seq analysis section. Peaks were called on merged replicates using MACS2 v2.2^90^. Bigwigs were z-score normalized as described above.

### CUT&Tag

Embryos were collected for one hour and aged for another two hours (2-3 h AEL). They were then dechorionated, rinsed in embryo wash buffer (PBS, 0.1% Triton X-100), and crude nuclear extracts prepared using a glass douncer in Nuclear Extraction buffer (20 mM HEPES pH 7.9, 10 mM KCl, 0.5 mM spermidine, 0.1% Triton X-100, 20% glycerol with protease inhibitor cocktail (Roche))^98^ and centrifuged at 700g for 10 min at 4°C. The nuclear pellets were resuspended in a Nuclear Extraction buffer and mixed with *Drosophila virilis* nuclei extracted from 3rd instar larvae (approximately 10% of the total number of nuclei). Nuclei corresponding to at least 100 embryos per reaction were incubated with 25 μl of BioMag®Plus Concanavalin A beads (Polysciences) (prepared in Binding buffer (20 mM HEPES pH 7.5, 10 mM KCl, 1mM CaCl2, and 1 mM MnCl2)) on a nutator for 10 min at 4°C. CUT&Tag was performed according to a published protocol^99^. Nuclei-bead complexes were resuspended in 100 μl antibody buffer (20 mM HEPES pH 7.5, 150 mM NaCl, 0.5 mM spermidine, 0.05% digitonin, 2 mM EDTA pH 8.0, and 0.1% BSA supplemented with protease inhibitor cocktail (Roche)). Antibodies were added and samples were incubated overnight at 4°C. We used the following antibodies: 1 μl rabbit H3K27ac (Abcam, ab4729), 1 μl rabbit H3K18ac (Abcam, ab1191), 1 μl rabbit H3K9ac (Abcam, ab4441), 1.2 μl rabbit CBP^58^, 1 μl TBP (a kind gift from Jim Kadonaga, UCSD), 1.2 μl rabbit anti-BRD4/fs(1)h (a gift of Renato Paro, kindly provided by Nicola Iovino^100^, 1.2 μl rabbit anti-CycT (a kind gift of Kazuko Hanyu-Nakamura)^101^, rabbit anti-RNA Polymerase II CTD repeat YSPTSPS (phospho-serine 5) (5SerP) (Abcam, ab5131), rabbit anti-RNA Polymerase II CTD repeat YSPTSPS (phospho-serine 2) (2SerP) (Abcam, ab5095). Following overnight incubation, the experimental procedure was followed using guinea pig α-rabbit antibody (Antibodies online cat. no. ABIN101961) secondary antibodies and purified pA-Tn5 (Protein Science Facility, KI, Stockholm). Tagmented DNA was PCR amplified using custom i5 and i7 PCR primers and Phusion® High-Fidelity PCR Master Mix with GC Buffer (NEB). PCR conditions were as follows: 72°C for 5 min, 98°C for 30 s, followed by thermocycling (98°C for 10 s and 63°C for 10 s) for 14 cycles and final extension at 72°C for 1 min. Amplified libraries were purified using Agencourt AMPure XP beads (Beckman Coulter) (1.1:1 bead to sample volume ratio). Libraries were paired-end (2 × 37 bp) sequenced on an Illumina NextSeq 2000 platform at the BEA core facility, Karolinska Institutet, Stockholm.

### CUT&Tag analysis

Sequencing data were uploaded to the Galaxy public server usegalaxy.org^102^. CUT&Tag reads with trimmed adapters were mapped to the *Drosophila melanogaster* (dm6) genome assembly using Bowtie2 (v.2.4.5) in the very sensitive local alignment mode. The unstranded BedGraph spike-in normalized files from individual replicates were generated using the BedTools (v.2.30.0) tool “Genome Coverage” using the default parameters^91^. A spike-in scaling factor was calculated as 10^8/[mapped *D. virilis* reads]. BedGraph files were converted to bigWig format, and then replicates were merged with the average signal by deepTools tool “bigwigCompare”^92^. CBP peaks were called using MACS2 (v. 2.2.7.1) with the following parameters: --mfold 30, 100 --bw 400 --qvalue 1e-7. CycT and Brd4 peaks were called with the following parameters: --mfold 5,50 --bw 300 --qvalue 0,0001.

### Precision run-on sequencing (qPRO-seq)

A variant of PRO-seq, qPRO-seq was performed on CBP^deGrad^ and CBP^CRY2^ embryos collected for one hour and aged for a further two hours (2–3 h AEL). Collected embryos were dechorionated in dilute bleach and rinsed thoroughly in embryo wash buffer (PBS, 0.1% Triton X-100) before being flash-frozen in liquid nitrogen and stored at −80°C. qPRO-seq was performed as previously described^103,104^. Briefly, embryos were resuspended in cold nuclear extraction buffer A (10 mM Tris-HCl pH 7.5, 300 mM sucrose, 10 mM NaCl, 3 mM CaCl2, 2 mM MgCl2, 0.1% Triton X, 0.5 mM DTT, protease inhibitor cocktail (Roche) and 4 µ/ml RNase inhibitor (SUPERaseIN, Ambion)), transferred to a dounce homogenizer and dounced with the loose pestle for 20 strokes. To remove large debris, the suspension was passed through Miracloth tissue (Merck, 475855-1R) followed by douncing with a tight pestle for 10 strokes. Nuclei were pelleted at 700g for 10 min at 4°C and washed twice in buffer A and once in buffer D (10 mM Tris-HCl pH 8, 25% glycerol, 5mM MgAc2, 0.1 mM EDTA, 0.5 mM DTT). For qPRO-seq, we used 2 million nuclei resuspended in buffer D and stored at −80°C. For further spike-in normalization, they were mixed with *Drosophila virilis* embryo nuclei (2% of the total number of nuclei). Nuclear run-on assays were performed in biological duplicates exactly as previously described using all four biotinylated dNTPs^103,104^. qPRO-seq libraries were sequenced (single-end 1 × 75 bp) on the Illumina NextSeq 2000 platform at the BEA core facility, Karolinska Institutet, Stockholm.

### qPRO-seq analysis

Sequencing data were uploaded to the Galaxy public server usegalaxy.org^102^. qPRO-seq reads were mapped after removing adapters to the *Drosophila melanogaster* (dm6) genome assembly using Bowtie2 (v.2.4.5) with the very sensitive end-to-end analysis mode^89^. The bam files were deduplicated using UMI, and the strand-separated spike-in normalized BedGraph coverage files from individual replicates were generated using the BedTools (v.2.30.0) tool “Genome Coverage” using the default parameters^91^. Strand-separated BedGraph were converted to bigWig format, and then two replicates of the same strand were merged with the average signal by deepTools tool “bigwigCompare”^92^.

To identify genes with differential nascent transcription, we used DEseq2 with estimateSizeFactor provided by user that equals to spike-in scaling factor. DEseq2 compared length-normalized read counts obtained by featureCounts on the bodies of shortest transcripts for each gene (defined as 500 bp downstream of the TSS to 100 bp upstream of the TES). The statistically significant difference was defined as padj < 0.05 and |log2 fold change| > 1. Genes in the first quartile by the number of reads mapping to the gene body were removed from the analysis. Genes with normalized counts below 50 for CBP^CRY2^ samples and 20 for CBP^deGrad^ samples were rejected. To examine Pol II promoter-proximal pausing, the promoter read counts (defined as 50 bp upstream of the TSS to 100 bp downstream of the TSS) were extracted with featureCounts and compared by DEseq2.

