## Supplemental Figures for "Catalytic-dependent and independent functions of the histone acetyltransferase CBP promote pioneer factor-mediated zygotic genome activation"

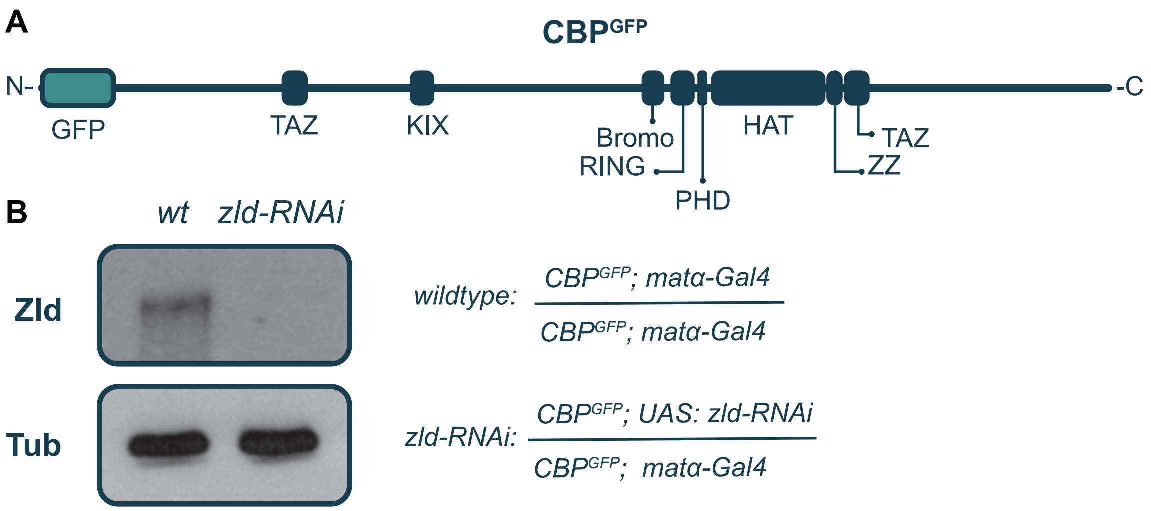


**C**

### Top enriched motifs

| Peak Class | Rank | Motif | Factor | p-value |
|---|---|---|---|---|
| Shared | 1 | GCAGGTAG | Zld | 4.7E-27 |
| | 2 | BDDBGCAGGTAGNND | Zld | 2.1E-20 |
| | 3 | ACCT | Zld | 1E-19 |
| Zld-Unique | 1 | GCAGGTAG | Zld | 2E-114 |
| | 2 | BDDBGCAGGTAGNND | Zld | 3.5E-83 |
| | 3 | DDBGCAGGTAGN | Zld | 4.6E-81 |
| CBP-Unique | 1 | TATCGATA | BEAF-32 | 1.8E-91 |
| | 2 | WNWTATCGATA | Dref | 5E-61 |
| | 3 | ATAWCGATAR | Dref | 1.2E-54 |

### Enrichment of Dl, Bcd, Twi motifs

| Peak Class | Motif | Factor | p-value |
|---|---|---|---|
| Shared | GGGAAAAYCC | Dl | 2.9E-07 |
| Zld-Unique | BGGRWTTYCCC | Dl | 0.00024 |
| CBP-Unique | GGGAAAAYCC | Dl | 0.00583 |
| Shared | VRGATTAD | Bcd | 0.00156 |
| Zld-Unique | SYTAATCC | Bcd | 0.00027 |
| CBP-Unique | RCCAGATGTB | Twi | 4.3E-06 |

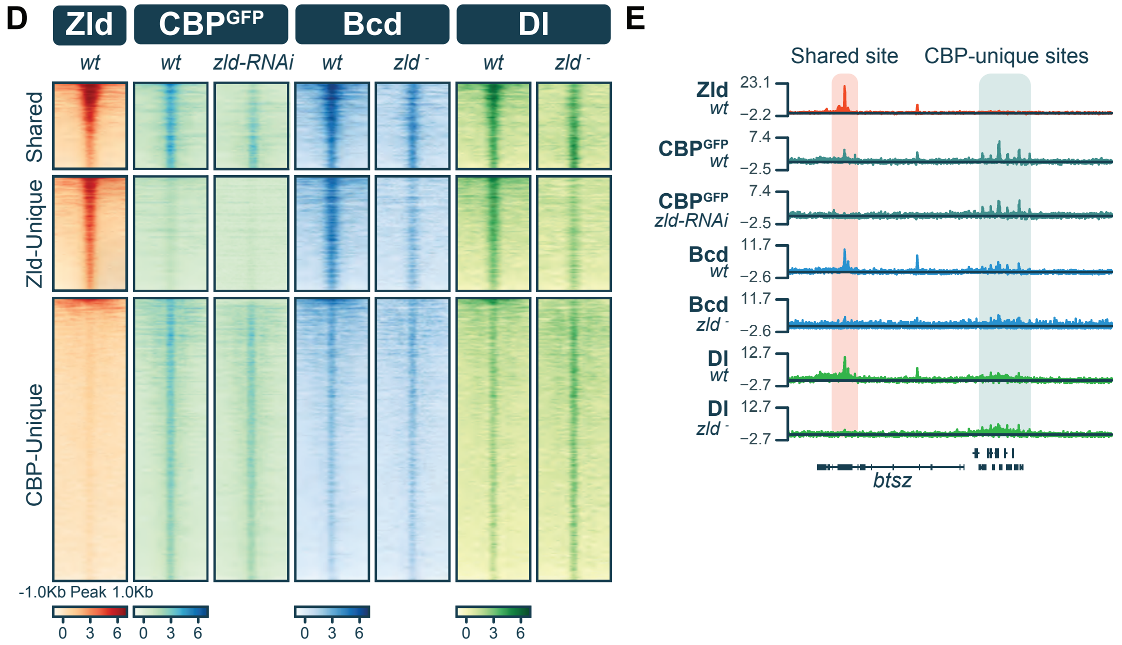


**Figure S1. Dl- and Bcd-binding motifs are enriched at CBP-bound ChIP-seq classes.**
**(A)** Model depicting protein structure of endogenous CBP$^{GFP}$ with predicted protein domains from UniProt database. **(B)** Immunoblot for Zld on extract from stage 5 wild-type and *zld-RNAi* embryos. Genotypes are as indicated. Tubulin is shown as a loading control. **(C)** Motif enrichment within peak classes identified by ChIP-seq. The top three enriched motifs of each class (left) and enrichment in each peak class for motifs bound by Zld-pioneer dependent transcription factors Dorsal (Dl), Bicoid (Bcd), and Twist (Twi) (right) are shown. **(D)** Heatmaps for ChIP-seq for Zld, CBP, Bcd and Dl in wild-type or *zld-RNAi* background as indicated. **(E)** Representative genome browser tracks of the ChIP-seq data shown in **(D)**.

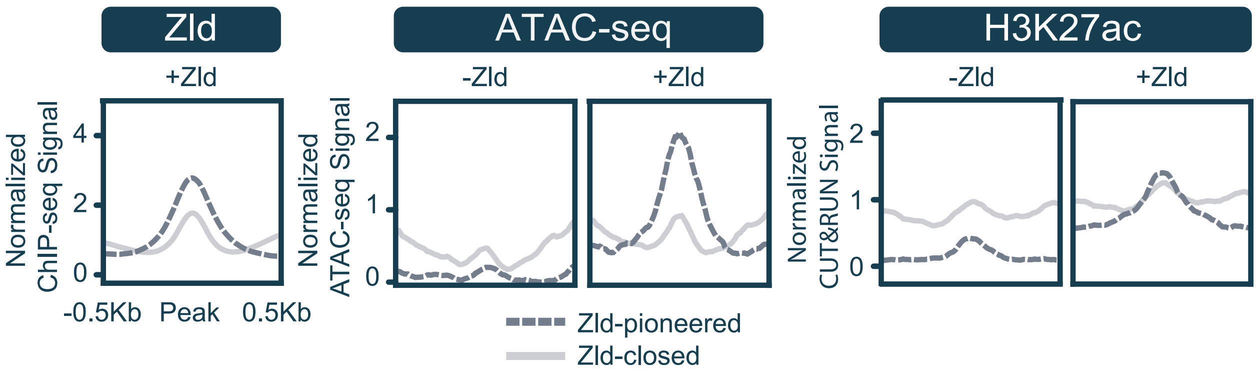


**Figure S2. H3K27ac is induced at Zld-pioneered regions in S2 cells.**
Metaplots of the average Zld occupancy (ChIP-seq), chromatin accessibility (ATAC-seq), or H3K27ac levels (CUT&RUN) from S2 cells. Data is z-score normalized and split into Zld-pioneered (dashed line) and Zld-closed (solid line) classes.

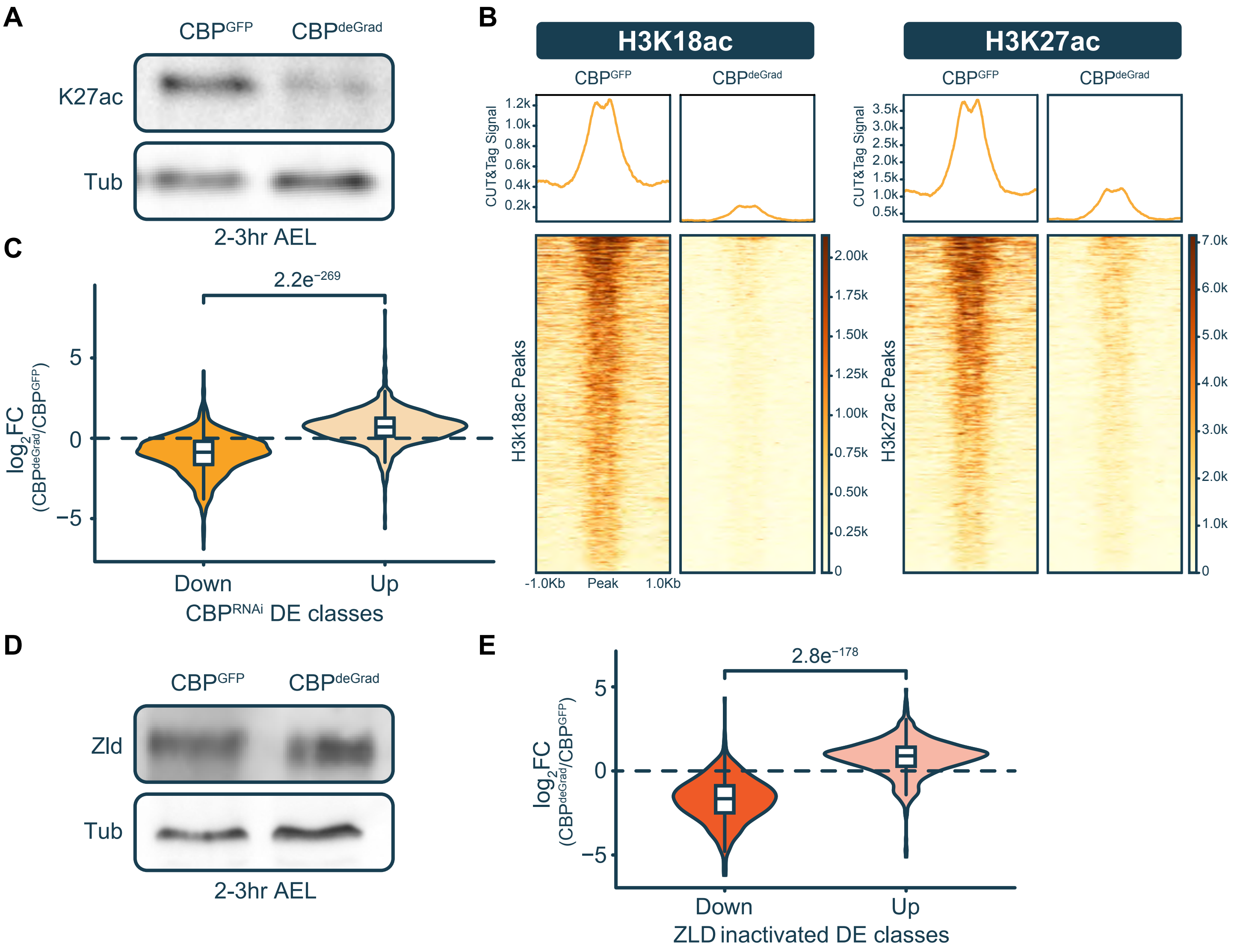


**Figure S3. CBP does not affect Zld expression and is required to activate Zld-dependent genes. (A)** Immunoblot for H3K27ac on extract from CBP$^{GFP}$ and CBP$^{deGrad}$ embryos 2-3hrs AEL. Tubulin is shown as a loading control. **(B)** Heatmaps and metaplots of CUT&Tag for H3K18ac and H3K27ac on CBP$^{GFP}$ and CBP$^{deGrad}$ embryos 2-3hrs AEL. **(C)** Violin plot of $\log_2$ fold change in gene expression in CBP$^{deGrad}$ as compared to CBP$^{GFP}$ embryo for genes differentially expressed (DE) in CBP$^{RNAi}$ embryos as compared to wild type (increased and decreased gene sets are indicated below) (adjusted p-value < 0.05, $|\log_2$ fold change$| > 1$) (data for CBP$^{RNAi}$ embryos are from Ciabrelli et al. 2023[30]). Statistics were calculated using a Wilcoxon Rank Sum test. **(D)** Immunoblot for Zld on extract from CBP$^{GFP}$ and CBP$^{deGrad}$ embryos 2-3hrs AEL. Tubulin is shown as a loading control. **(E)** Violin plot of $\log_2$ fold change in gene expression in CBP$^{deGrad}$ as compared to CBP$^{GFP}$ embryo for genes differentially expressed (DE) in Zld-inactivated embryos as compared to untreated embryos (increased and decreased gene sets are indicated below) (adjusted p-value < 0.05, $|\log_2$ fold change$| > 1$) (data for Zld-inactivated embryos are from McDaniel et al. 2019[15]). Statistics were calculated using a Wilcoxon Rank Sum test.

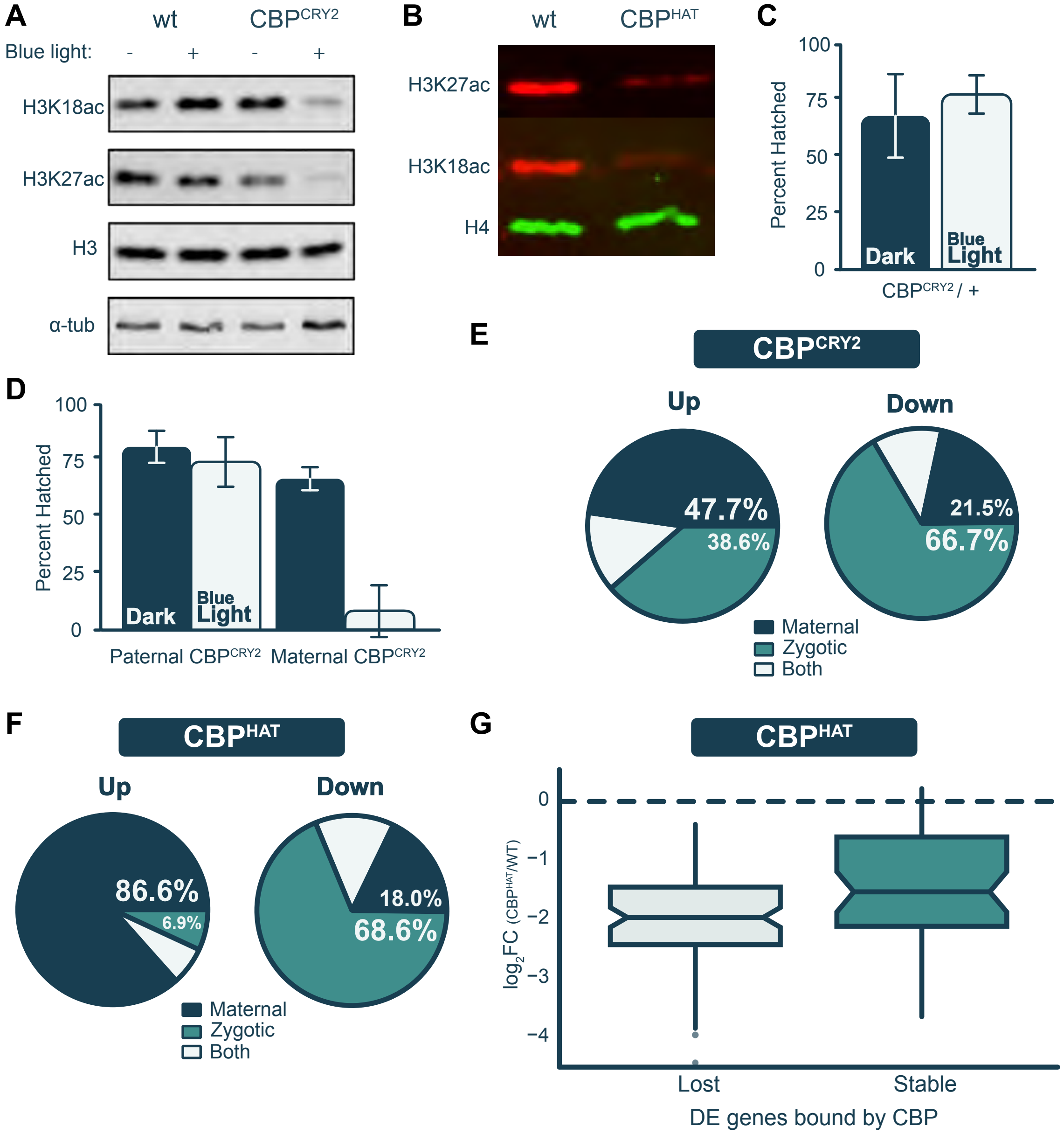


**Figure S4. Maternally encoded, catalytically active CBP is required for histone acetylation, embryo viability and zygotic transcription.**

**(A)** Immunoblots for H3K18ac, H3K27ac, and histone H3 on extract from stage 5 wild-type (wt) or CBP$^{CRY2}$ embryos treated with (+) or without blue light (-) during the MZT. Tubulin is shown as a loading control. **(B)** Immunoblots for H3K18ac, H3K27ac, and histone H4 on extract from wild-type (wt) and CBP$^{HAT}$ stage 5 embryos. **(C)** Hatching rates for progeny from heterozygous CBP$^{CRY2}$ mothers (n=300 both blue-light treated and raised in the dark). **(D)** Hatching rates of heterozygous embryos that inherited either a paternal or maternal copy of *CBP$^{CRY2}$* that were raised in the dark (paternal n= 217, maternal n=310) or treated with blue light (paternal n= 304, maternal n=301) during the MZT (0-3hrs AEL). Error bars are the standard deviation between replicates. **(E-F)** Pie charts indicating the timing of gene expression during the MZT (maternal, zygotic, or both) for the genes with increased (left) or decreased (right) expression in CBP$^{CRY2}$ embryos treated with blue light as compared to those in the dark **(E)** or CBP$^{HAT}$ embryos compared to wild type **(F)**. **(G)** Box plot showing the differentially expressed genes in CBP$^{HAT}$ embryos as compared to controls that are bound by CBP in wild-type, control embryos. These genes are classified by those that lose CBP occupancy in CBP$^{HAT}$ mutants (Lost) or maintain CBP occupancy (Stable).

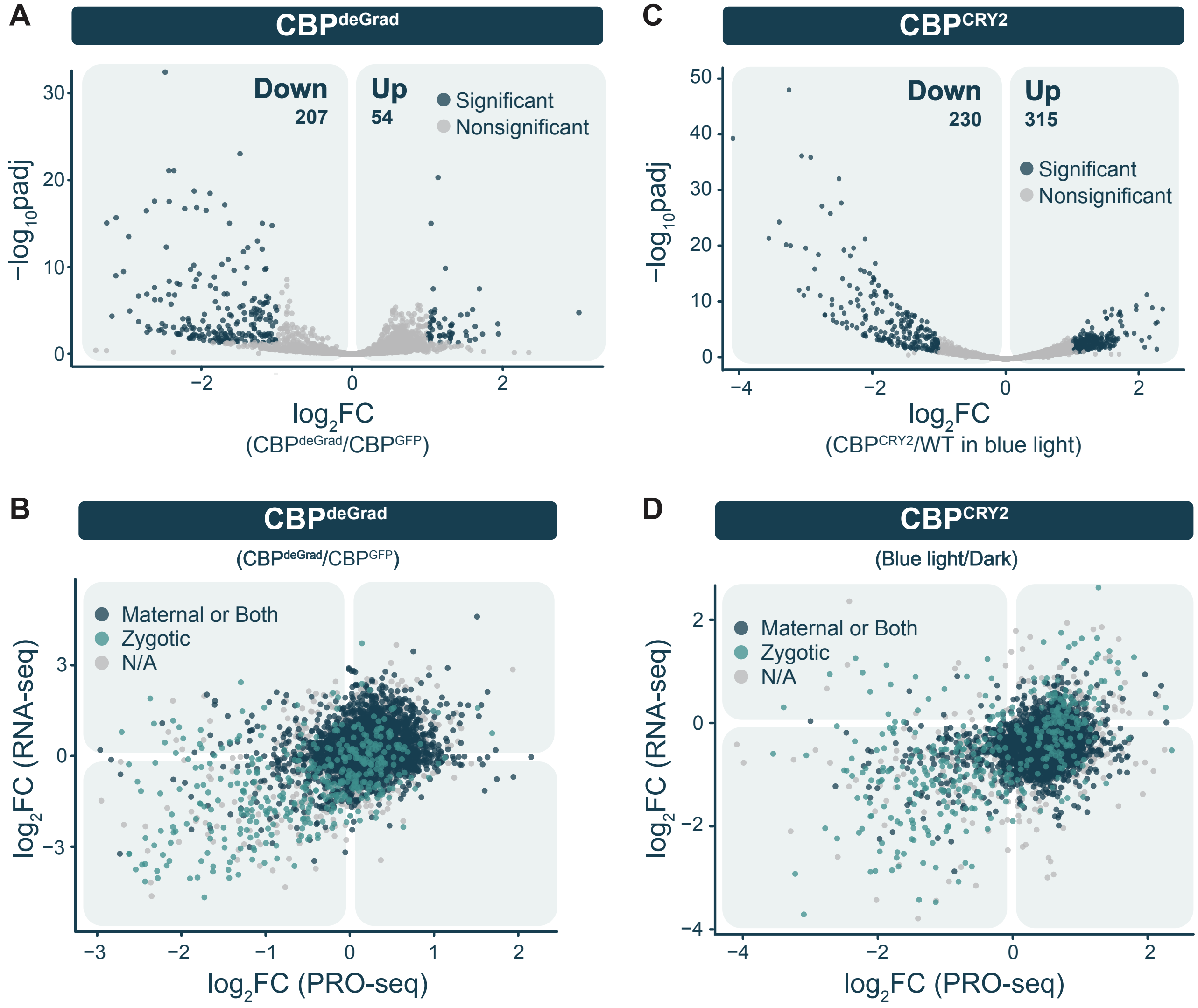


**Figure S5. CBP is required for nascent transcription in the early embryo.**
**(A,C)** Volcano plots of gene body PRO-seq from $CBP^{deGrad}$ embryos as compared to $CBP^{GFP}$ embryos **(A)** and $CBP^{CRY2}$ embryos treated in the dark (-) or in blue light (+) **(C)**. **(B,D)** Scatter plots comparing PRO-seq with RNA-seq changes in $CBP^{deGrad}$ embryos as compared to $CBP^{GFP}$ embryos **(B)** and $CBP^{CRY2}$ embryos treated in the dark (-) or in blue light (+) **(D)**.

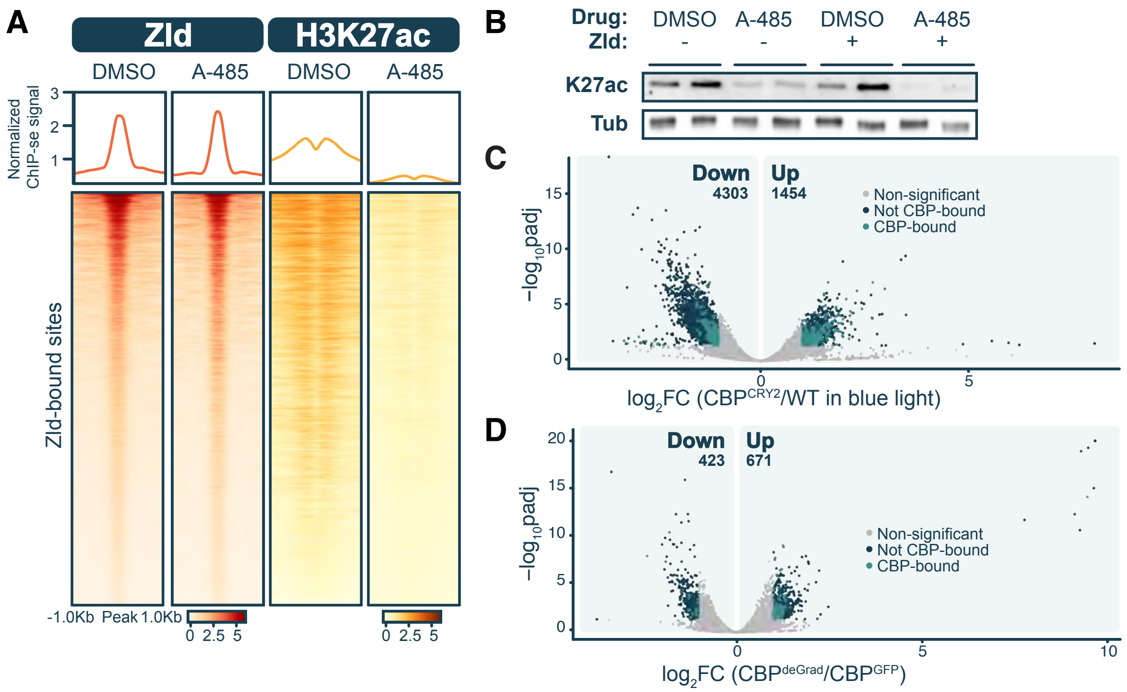


**Figure S6. CBP is not required for accessibility at Zld-pioneered sites or in the early embryo.** **(A)** Heatmaps of Zld occupancy or H3K27ac levels (ChIP-seq) at Zld-bound regions in S2 cells treated with DMSO or A-485. Zld ChIP-seq are z-score normalized. H3K27ac ChIPseq are spike-in normalized. **(B)** Immunoblot for H3K27ac on S2 cell extracts. Zld expression was induced with CuSO4 at the concentrations indicated and treated with DMSO or A-485. Tubulin is shown as a loading control. **(C-D)** Volcano plots of single-embryo ATACseq peaks from $CBP^{CRY2}$ embryos compared to wild-type embryos treated with blue light **(C)** or $CBP^{deGrad}$ embryos as compared to $CBP^{GFP}$ embryos **(D)**. Regions bound by CBP as identified in ChIP-seq on $CBP^{GFP}$ embryos are indicated in teal. Navy represents genes that change in expression that lack proximal CBP-binding sites, and grey represents genes with statistically insignificant changes in expression (adjusted p-value < 0.05, |log2 fold change| > 1).
